# Are dogs with congenital hearing and/or vision impairments so different from sensory normal dogs? A survey of demographics, morphology, health, behaviour, communication, and activities

**DOI:** 10.1101/2020.03.06.980482

**Authors:** Sophie Savel, Patty Sombé

**Affiliations:** Aix Marseille University, CNRS, Centrale Marseille, LMA UMR 7031, Marseille, France; Non-profit organisation “Blanc Comme Neige”, Saint-Denis-sur-Scie, France

## Abstract

The births of domestic dogs with pigment deletion and associated congenital hearing and/or vision impairments are increasing, as a result of mutations of certain genes expressing popular coat colour patterns (Merle, piebald, Irish spotting). The future of these dogs is often pessimistic (early euthanasia or placement in rescues/fosters, lack of interactions and activities for adults). These pessimistic scenarios result from popular assumptions predicting that dogs with congenital hearing/vision impairments exhibit severe Merle-related health troubles (cardiac, skeletal, neurological), impairment-related behavioural troubles (aggressiveness, anxiety), and poor capacities to communicate, to be trained, and to be engaged in leisure or work activities. However, there is no direct scientific testing, and hence no evidence or refutation, of these assumptions. We therefore addressed an online questionnaire to owners of 223 congenitally sensory impaired (23 vision impaired, 63 hearing impaired, 137 hearing and vision impaired) and 217 sensory normal dogs from various countries. The sensory normal cohort was matched in age, lifetime with owner, breed and sex with the sensory impaired cohort, and was used as a baseline. The questionnaire assessed demographics, morphology, sensory impairments, health and behavioural troubles, activities, and dog-owner communication. Most hearing and vision impaired dogs exhibited abnormal pigment deletion in their coat and irises. Vision impaired dogs additionally exhibited ophthalmic abnormalities related to Merle. The results refute all above-listed assumptions, except for neurological troubles. We however suggest that reports of neurological troubles could be partially accounted for by lacks of diagnosis of breed-related drug sensitivity and impairment-related compulsive behaviours. Results about communication and activities are particularly optimistic. The need for future studies of numerous dogs from various breeds tested for Merle, piebald and medical-drug-resistance genes, and the beneficial effects that present and future research may have on the future of sensory impaired dogs, are discussed.

## Introduction

In order to meet an increasing demand for pet dogs, most countries report growing numbers of dog breeders, atypical phenotypes in either existing or novel dog breeds, and births of puppies with various genetic defects [1]. For example, the population of dogs with congenital hearing and/or vision impairments is increasing. This population, which is the focus of the present study, mostly results from mutations of the genes that express certain of the most popular coat colour patterns.

### Demographics and genetics of congenitally sensory impaired dogs

One of the most popular coat patterns in dogs is Merle. The Merle coat can be described as a patchwork randomly composed of areas of full pigmentation combined with areas of lighter, diluted pigmentation. Originally, the Merle trait was essentially produced in certain breeds, mainly from the herding group. This trait has progressively been introduced in a growing number of, sometimes unexpected, breeds. To date, Merle can be found in the following breeds, listed in alphabetic order: Alapaha Blue Blood Bulldog, American Cocker Spaniel, American Pit Bull Terrier, American Staffordshire Terrier, Australian Koolie, Australian Shepherd, Border Collie, Boxer, Chihuahua, Collie, Dachshund, French Beauceron, French Bulldog, Great Dane, Hungarian Mudi, Labradoodle, Louisiana Catahoula, Lurcher, Miniature American Shepherd, Norwegian Dunkerhound, Schnauzer, Shetland Sheepdog, Pomeranian, Poodle, Pyrenean Shepherd, and Welsh Sheepdog [2–4]. The Merle coat is particularly frequent in Australian Shepherds, followed by Border Collies, Great Danes, and Shetland Sheepdogs. To note, Merle is not accepted for registration in kennel clubs for several of the breeds listed.

The Merle coat is expressed by the gene of same name located on the M locus, that is inherited in an autosomal, incomplete dominant manner [5]. Merle is, at the homozygous “double Merle” state, one the four known pigment genes in dogs, along with piebald [2], Irish spotting [2] and KIT [6], whose mutation has the deleterious effect of deleting pigments in hairs, skin, nose and mucous, iris and tapetum lucidum, and stria vascularis of the inner ear. The lack of pigments in the stria vascularis causes early death of sensory hair cells in the scala media. As a result, dogs with mutated above-mentioned genes have excessive white coat, pink skin, nose and mucous, light blue irises, and congenital, sensorineural, irreversible hearing impairments.

As for Merle, piebald concerns various breeds, but this trait, located on the S locus, is recessive. Pigment deletion and hearing impairments mostly occur in homozygous piebalds. No genetic testing is yet available for Irish spotting, although this gene is assumed to be present in numerous breeds. The KIT mutation is less statistically problematic, because it exclusively concerns the German Shepherd breed and is early embryonic lethal at the homozygous state.

Contrary to piebald, Irish spotting and KIT, Merle is additionally associated with various congenital ophthalmic abnormalities, referred to as Merle Ocular Dysgenesis. The most severe ophthalmic abnormalities are observed in homozygous Merles [7–8]. Ophthalmic abnormalities in homozygous Merles can concern the eyeball (reduced size, called microphthalmia, or total absence), the cornea (microcornea), the iris (coloboma, hypoplasia), the size, shape, position or reaction of the pupil (starburst or misshapen pupil, dyscoria, corectopia), the pupillary membrane (persistence), the lens (cataract, microphakia, luxation), the sclera (coloboma, staphyloma), the retina (detachment, dysplasia), and the optic nerve. Depending on the severity of their ophthalmic abnormalities, homozygous Merles can exhibit moderate to severe vision impairments. These vision impairments related to Merle are susceptible to worsen, or even to appear, over the life course.

It has long been assumed that the M locus has two possible alleles, namely non-merle (m, expressing solid phenotype) and Merle (M, expressing Merle phenotype). Latest research has identified between four and six variations of the Merle allele, as a function of the length of the poly-A tail of its SINE insertion [**9-10; 3**]. The most detailed work is that conducted by Langevin and colleagues, who have tested hundreds of dogs from various breeds on the M locus and have determined six Merle allele variations [3–4]. Their goals were to accurately determine which SINE insertion lengths can express a Merle pattern, which phenotype is most typically associated with each possible genotype, which cases of mosaicism can occur, and which breeding between genotypes are susceptible to produce excess white, sensory impaired, double Merle puppies. However, these genetics studies of Merle are very recent. The state-of-the-art equipment needed for precise examination of SINE insertion length and mosaicism is recent and expensive. Therefore, only two of the 16 laboratories that propose Merle testing in dogs can currently provide this detailed information in their test results. Few of the remaining laboratories provide state-of-the-art information about Merle genetics on their public websites [11]. As a result, many dog breeders and owners are not fully aware of the complexity of Merle genetics and the conditions of at-risk breeding. Many countries have not yet strictly regulated Merle breeding. Comparable lacks of information and regulation are observed for different breeds with piebald trait.

For all the reasons mentioned above, births of excess white puppies with congenital, sensorineural, irreversible hearing and/or vision impairments are still frequently observed worldwide. There are different possible scenarios regarding the future of these puppies. Non-professional breeders and private individuals with little knowledge of the genetics of sensory impairments sell their puppies as exotic specimens without providing any adequate information to buyers. Numerous professional breeders with informed knowledge either have the puppies euthanized shortly after birth or entrust them after weaning to specialised rescue centres or foster programs that have very restrictive adoption criteria for such dogs. Dogs that have been lucky enough to be adopted and to become adults often live in controlled – sometimes “overprotective” – environments, and do not often have access to canine activities. All these pessimistic scenario result from a series of popular assumptions about deaf and/or blind dogs, that are detailed below.

### Assumptions about sensory impaired dogs

It is often assumed that excess white dogs, particularly double Merles, exhibit severe, or even lethal, health issues in their neurological, cardiac, skeletal and reproductive systems. We propose below three possible origins of these possibly false assumptions.

First, the assumption of lethality in excess white, double Merle dogs may originate from the fact that, in horses, the mutation of the Overo gene causes both abnormal white coat and early death of the foal [12]. A popular belief has incorrectly extended this relatively well-known fact to all mammalian animal species. Accordingly, many websites and social media relative to canine genetics refer to excess white dogs as “lethal white dogs”. In fact, there are only four canine genes that have proved to be – early embryonic – lethal at the homozygous state (KIT in German Shepherds, “Natural Bobtail” in various breeds, Harlequin in Great Danes, and “Hairless” in Chinese, Mexican and Peruvian hairless breeds) [4]. Merle is not one of them.

Second, the assumption of neurological issues in doubles Merles may originate in part from the fact that the most common neurological disorder in dogs, namely primary idiopathic epilepsy, is frequent in certain breeds from the herding group, such as Australian Shepherds and Border Collies [13]. As specified above, these two breeds are most frequently concerned by the homozygous Merle genotype. The assumption of neurological issues may additionally, or even above all, result from the fact that Australian Shepherds and Border Collies are also frequently concerned by mutations of the medical drug resistance (MDR1) gene [14–15]. As detailed further below, mutated MDR1 gene elicits neurotoxic, sometimes epileptiform reactions to common chemical agents (*e.g*., parasite control products) that are well tolerated by dogs with normal MDR1 gene. In other words, whether the neurological signs observed in double Merle Australian Shepherds and Border Collies are linked to Merle, as frequently assumed, or to breed-dependent disorders and MDR1 mutation, is undetermined.

Third, assumptions of cardiac, skeletal and reproductive issues in doubles Merles may essentially originate from multiple citations of a single study, that just contained the following short statement in the Introduction: “*In all breeds, the double merle genotype can be sublethal and is associated with multiple abnormalities of the skeletal, cardiac, and reproductive systems*” [5]. However, this study was on genetic testing of Merle, not on health, and cited three studies to support the statement [16–18]. These three studies were conducted long before genetic testing of Merle was available, examined either small or poorly genetically diversified dog cohorts (*i.e*.; total of 32 dogs or single genealogical branch), and never clearly referred to the types of health issues that are nonetheless listed in the statement quoted above.

Moreover, it is often assumed that congenitally deaf and/or blind dogs exhibit behavioural troubles (see foreword by Strain in [19]). Principally, their severe sensory impairment(s) are believed to increase frustration, and to elicit resultant aggressiveness and anxiety troubles. Also, it is assumed that deaf and/or blind dogs are particularly susceptible to bite because they are easily startled when they are approached. Abnormal brain structures, and concomitant abnormal mental capacities, have also been assumed in congenitally deaf dogs. However, this assumption is based on a single neuro-imagery study that just reported a reduction in size of the auditory cortex in congenitally deaf Dalmatians [20].

For dogs as for many social species, hearing and vision are two important sensory modalities for conspecific and interspecific communication [21]. Thus, deaf and/or blind dogs are believed to have poorer communication capacities, in particular with their human caregivers. As a result, it is often assumed that deaf and/or blind dogs cannot be trained, and cannot be safely and efficaciously engaged in any – individual or collective, conspecific or interspecific, leisure or work – activity.

### Aims and methodological choices of the study

In summary, above-mentioned assumptions predict that congenitally hearing and/or vision impaired dogs frequently exhibit health and behavioural troubles, and are poorly capable of communicating and practicing activities. These assumptions are so popular that they have drastic consequences on the future of sensory impaired puppies. However, there is to date no scientific evidence or refutation of these assumptions. Precisely, there is no study that we are aware of that directly assessed either health, behaviour, communication or activities in congenitally sensory impaired dogs, or that compared sensory impaired and sensory normal dogs on these points. One exception is the study by Farmer-Dougan and colleagues, who addressed a survey of behavioural traits to owners of hearing/vision impaired and sensory normal dogs [22]. They found lower scores of aggressiveness and anxiety in the former cohort, which is opposite to the assumption.

The aims of the present study were therefore to examine health, behaviour, communication and activities for a cohort of congenitally hearing and/or vision impaired dogs, and to compare the results with those from a “baseline” cohort of sensory normal dogs that was matched in breed, age, sex and lifetime with owner with the sensory impaired cohort. Additionally, we aimed to gain insight concerning the diagnosis of sensory, particularly hearing, impairments. The sole way to assess unilateral hearing impairment in dogs is to conduct objective measurements of brainstem auditory evoked responses (BAER). However, animal BAER testing sites are infrequent (*e.g*., see list in [19]). Little is known about how exactly hearing is subjectively evaluated by veterinaries, breeders, owners, *etc*, in the numerous dogs that have no access to BAER testing. Finally, we aimed to verify whether congenital sensory impairments were frequently associated with pigment deletion in the coat and irises and with ophthalmic abnormalities. As such, the genetic cause of the sensory impairments was indirectly explored.

To assess these different points, we chose to conduct an owner survey, a method that is frequently employed to assess, for example, health [23] and behaviour [22], in dogs. Surveys of dog owners have some disadvantages relative to the degrees of interest, understanding, recall and impartiality of the respondents, but have above all multiple advantages. They allow the inclusion of dogs with much wider characteristics compared to surveys of veterinaries or animal insurances for questions on health, dog behaviourists for questions on behaviour, or canine clubs for questions on activities. In other words, owner surveys are not restricted to dogs *with* health/behavioural issues and activities. We chose the online diffusion of the questionnaire in two languages, namely French and English, in order to expand its geographic distribution.

## Materials and methods

### Questionnaire content

The questionnaire contained 30 questions about dogs, divided into 7 sections:

- **Demographics**: country; date of birth; date of acquisition; site of acquisition; sex; breed; presence of other dog(s) at home.
- **Morphology**: surface of white coat on body and head; colour of irises; ophthalmic abnormalities; for sensory impaired dogs only: is there indication, from either genetic testing or parental phenotypes, that the dog is double Merle?
- **Determination of sensory impairments**: sensory status (*i.e*., normal, partially impaired, totally impaired) at each ear and each eye; type of diagnosis test (*i.e*., objective or subjective); operator of the subjective test; for hearing impairments only: stimuli and conditions of the subjective test.
- **Health**: has the dog ever suffered from neurological, heart, bones/joints, skin, digestive or other health troubles? has the dog been tested for the MDR1 gene? if so, did the test indicate MDR1 mutation, and hence abnormal drug sensitivity?
- **Behaviour**: has the dog ever suffered from aggressiveness, anxiety, attention deficit/hyperactivity disorder (ADHD), obsessive compulsive disorder (OCD), or other behavioural troubles? who diagnosed the behavioural trouble(s)? have drugs been prescribed for this/these trouble(s)?
- **Activities**: frequency of practice of a series of leisure/sport activities; level at which each reported activity is practiced; is the dog engaged in assistance/therapy activities with either elderly, blind, or diabetic/epileptic persons?
- **Interspecific communication**: types of vocalisations produced by the dogs to communicate with their owners; types of signs used by the owners to communicate with, and train, their dogs.

All respondents gave their informed consent for the anonymous use and publication of their responses. They were proposed to send a picture of their dog to the first author by email. We received 88 pictures, that are presented in **S1 Fig** as illustrative examples of coat colour and ophthalmic abnormalities. The study was carried out in accordance with the ethical standards of the institutional review board at the Aix-Marseille University.

### Survey distribution

The questionnaire was published online using Google forms in two languages: French and English (see screenshot of the English online questionnaire in **S2 Fig**). Both versions were operational online from 19^th^ April 2019 until 30^th^ September 2019. Calls for participation, that included a short description of the survey and a direct link to the google form, were published on a variety of social media. The social media dealt with various canine themes, such as breeding, genetics, sensory impairments, training methods, activities, veterinary medicine, and behaviour. Calls for participation specified that the questionnaire was addressed to owners of dogs:

- aged between 9 months and 12 years
- with either no or congenital hearing and/or vision impairments
- that belonged to breeds for which the Merle coat is – frequently or occasionally – observed.

Dogs with acquired, late onset sensory impairments resulting from trauma, age, medication, surgery, *etc*, were explicitly excluded. Owners had the possibility to fill the questionnaire several consecutive times for different individual dogs.

### Size and geographic dispersion of the sample

Following data collection, responses to the French version of the questionnaire were translated in their English equivalent. Responses to French and English versions were then gathered. Overall, owners of 510 individual dogs completed the survey. The responses relative to 75 dogs were excluded from data analysis because mandatory questions were inadequately responded and/or one above-mentioned criterion was not fulfilled. For example, 33 dogs were aged less than 9 months, 24 were aged more than 12 years, and 20 were from “out-of-subject” breeds (*e.g*., Beagle, Dalmatian, Yorkshire, Golden Retriever, unidentified mix of breeds, *etc*.). The final sample therefore included 440 individual dogs, whose geographic dispersion between continent and countries is presented in **Table 1**. About 85% of the dogs were from either France or United States of America. The remaining dogs were spread between 15 countries.

**Table 1.** List of countries and corresponding number of dogs.

| Continent | Country | Number of dogs |
| --- | --- | --- |
| Europe | France | 240 |
|  | Belgium | 16 |
|  | Switzerland | 7 |
|  | Netherlands | 4 |
|  | United Kingdom | 4 |
|  | Germany | 3 |
|  | Finland | 2 |
|  | Italy | 2 |
|  | Spain | 1 |
|  | Slovakia | 1 |
| America | United States of America | 136 |
|  | Canada | 9 |
|  | Mexico | 2 |
|  | Brazil | 1 |
| Oceania | Australia | 5 |
|  | New Zealand | 3 |
| Africa | South Africa | 4 |

Continent and countries are sorted by descending number of dogs.

### Constitution of groups based on sensory status

Responses about the hearing and vision sensory status of the dog (*i.e*., normal, partially impaired, or totally impaired) were used to classify the 440 dogs in four groups:

- **HNVI (Hearing Normal Vision Impaired**) **= 23 dogs**: response “normal” for hearing at both ears, and response “partially/totally impaired” for vision at either one or both eyes
- **HIVN (Hearing Impaired Vision Normal) = 63 dogs**: response “partially/totally impaired” for hearing at either one or both ears, and response “normal” for vision at both eyes
- **HIVI (Hearing Impaired Vision Impaired) = 137 dogs**: response “partially/totally impaired” for hearing at either one or both ears and for vision at either one or both eyes
- **HNVN (Hearing Normal Vision Normal) = 217 dogs**: response “normal” for hearing at both ears and for vision at both eyes.

In summary, the study included 223 sensory impaired dogs, classified in three groups, and 217 sensory normal dogs. The first two sensory impaired groups had one – either hearing or vision – impairment, while the third sensory impaired group had both hearing and vision impairments. In the Figures and Tables below, the three sensory impaired groups were frequently gathered in a single “IMP” (impaired) cohort. The HNVN, sensory normal group was used as a baseline for comparison with the sensory impaired groups/cohort.

### Statistical analysis

Owners were asked to report the exact dates of birth and acquisition of their dog. These two dates were used to determine the dog’s age and lifetime with owner, respectively, in years, at the day of participation in the survey. The normal distributions of individual age and lifetime values for each group and for the IMP cohort were assessed using Shapiro-Wilk tests. All distributions differed significantly from normal (*p* < 0.05). Between-group comparisons in both age and lifetime with owner were therefore assessed using non-parametric, Kruskal-Wallis tests.

The frequency of each categorical response under study was obtained for each group by dividing the number of times the response was reported in the group by the total number of dogs in that group and then multiplying by 100. The response frequencies obtained are presented below either in Figures (ordinate) or in Tables. Chi^2^ tests for unpaired data were used to statistically assess two-by-two differences between groups in categorical responses. Each Chi^2^ test compared the raw numbers of reported/A and non-reported/B responses obtained in one group (*e.g*., numbers of “yes” and “no” responses, numbers of “male” and “female” responses, *etc.*) with those obtained in the other group. The list of comparisons assessed using Chi^2^ tests is provided in **Table 2**.

**Table 2.**
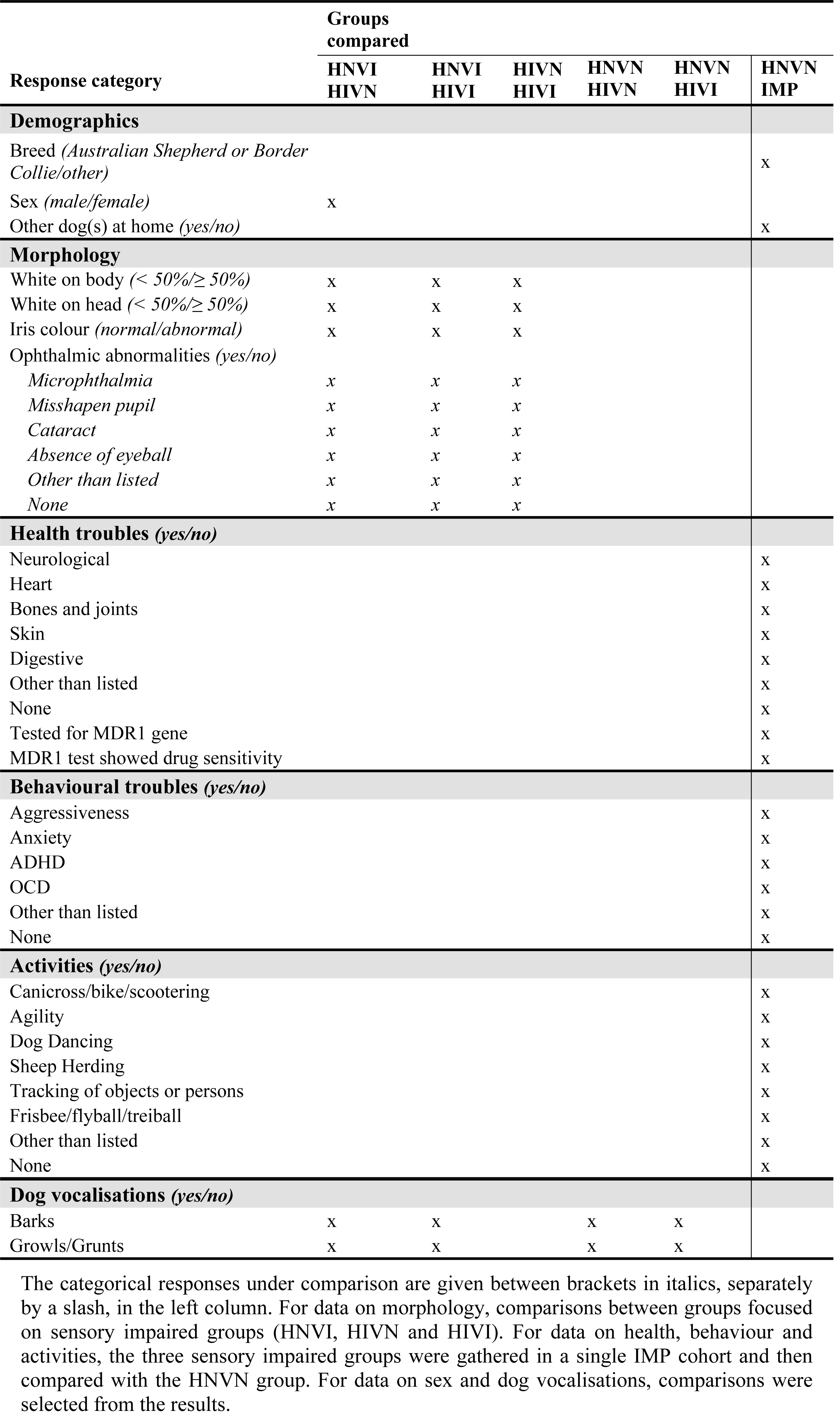
List of two-by-two comparisons between groups assessed using Chi^2^ tests.

Two-tailed *p* values reported in text and Figures were adjusted using the Holm’s correction for multiple comparisons. In Figures, *p* values for significant tests are emboldened and are followed by either four (*p* < 0.0001), three (*p* ≥ 0.0001 and < 0.001), two (*p* ≥ 0.001 and < 0.01) or one (*p* ≥ 0.01 and < 0.05) asterisk(s). *p* values for non-significant tests are reported in plain and are followed by “(ns)”.

## Results and discussion

### Demographics

#### Age

Box-plots of age values for each group and for the IMP cohort are presented in **Fig 1a**. There was no significant two-by-two difference in age between sensory impaired groups (median ages for HNVI, HIVN and HIVI groups = 2.6, 3.0 and 3.3 years, respectively; *H* < 0.25, *p* = 1.0) or between IMP and HNVN (median ages = 3.1 and 3.5 years, respectively; *H* = 3.87, *p* = 0.34). The five distributions had comparable lower quartiles (between 1.4 and 2.1 years), upper quartiles (between 5.0 and 5.8 years) and inter-quartile distances (between 3.1 and 3.8 years), and had few outlier values (frequencies ≤ 5%). Age has therefore unlikely contributed to between-group differences in the data examined in the different sections of the survey.

**Fig 1.**
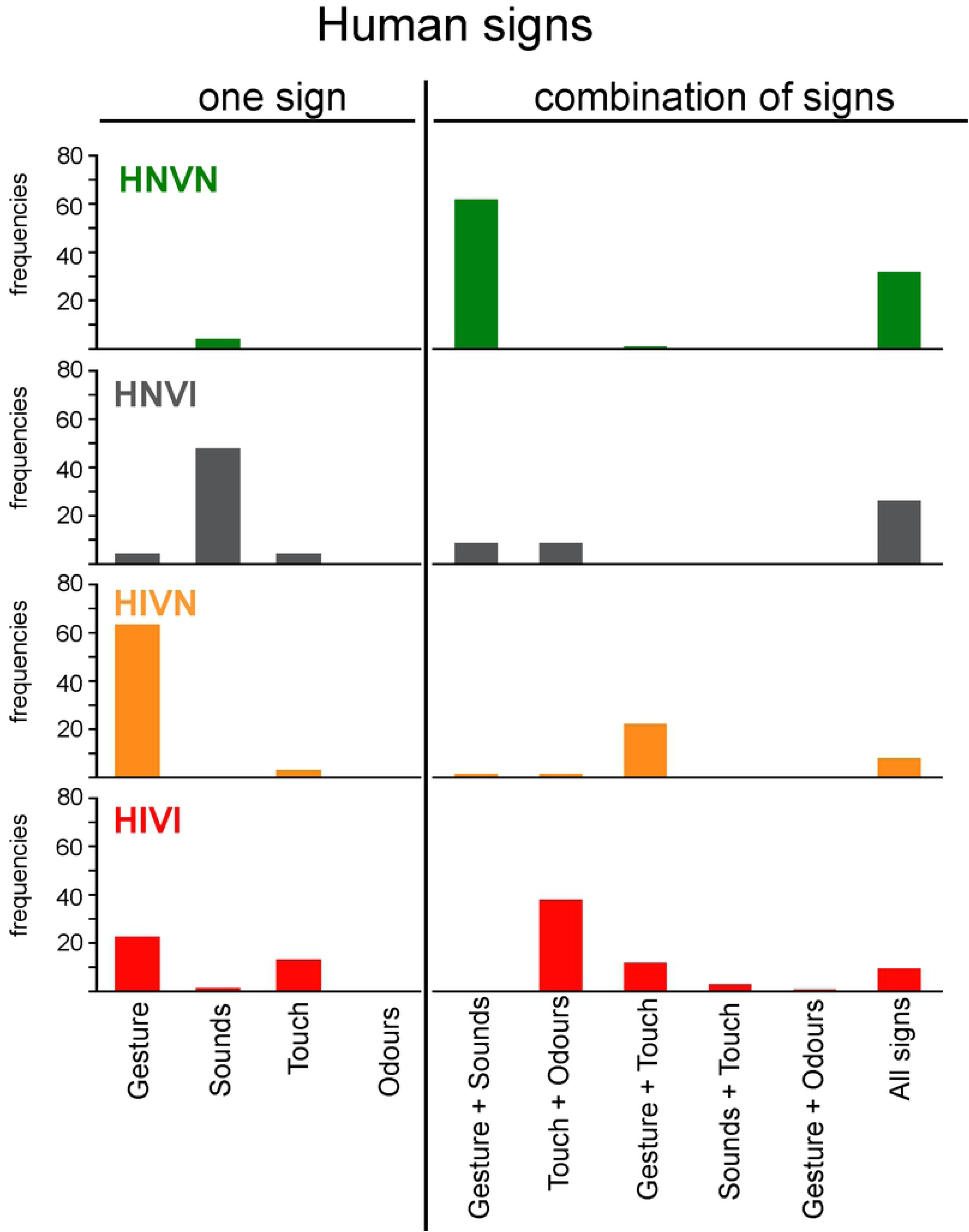
**Box-plots of individual values of (a) age and (b) lifetime with owner for each group, and for the three sensory impaired groups gathered (IMP).** HNVI = hearing normal vision impaired (grey), HIVN = hearing impaired vision normal (orange), HIVI = hearing impaired vision impaired (red), HNVN = hearing normal vision normal (green), IMP = impaired (purple), ns = not significant. The bold bar within the boxplot is the median, the cross is the mean, the bottom and top of the box are the lower and upper quartiles, respectively, and the dots are outlier values above the [upper quartile + 1.5 * inter-quartile distance] limit (top of upper vertical bars). Horizontal brackets indicate the two-by-two comparisons between groups that were statistically assessed using Shapiro-Wilk tests.

#### Lifetime with owner

Box-plots of lifetime values for each group and for the IMP cohort are presented in **Fig 1b**. There was no significant two-by-two difference between sensory impaired groups (median lifetimes with owner for HNVI, HIVN and HIVI groups = 2.2, 2.4 and 2.3 years, respectively; *H* < 0.24, *p* = 1.0), but the difference between IMP and HNVN cohorts was significant (median lifetimes with owner = 2.3 and 3.1 years, respectively; *H* = 13.50, *p* = 0.002). Both cohorts however had lower quartiles above one year. In behavioural studies of dog-owner communication that report the lifetime of the dyad, the smallest lifetime is one year [e.g., 24]. The difference in lifetime with owner between sensory impaired and sensory normal dogs in the present study have therefore unlikely contributed to differences between groups in the responses relative to interspecific activities and communication.

#### Breed

Owners were asked to report the breed of their dog using a list of purebred and mixed breeds followed by a field for manual report of non-listed breeds (see **S2 Fig**). The frequencies of the different responses obtained for each group are provided in **Table 3a**. The four groups were essentially composed of breeds from the herding group. Australian Shepherds and Border Collies, either purebred or mixed, represented 79% and 84%, respectively, of sensory impaired and sensory normal cohorts (*X^2^* = 0.22, *p* = 1.0, ns). Potential effects of breed on the data examined below may have therefore been equally elicited in all groups.

**Table 3.** Frequencies of responses, in percentages, obtained for each group and for the IMP cohort for the following demographic data: (a) breed, (b) sex, (c) site of acquisition, and (d) presence of other dog(s) at home.

| **Demographic data** | **HNVI** | **HIVN** | **HIVI** | **HNVN** | **IMP** |
| --- | --- | --- | --- | --- | --- |
| **(a) Breed** |  |  |  |  |  |
| **Herding group** |  |  |  |  |  |
| Australian Shepherd | 39 | 49 | 41 | **51** | **43** |
| Australian Shepherd x Border Collie | 4 | 8 | 9 | **3** | **8** |
| Australian Shepherd x unknown | 4 | 5 | 11 | **2** | **9** |
| Border Collie | 26 | 19 | 12 | **25** | **15** |
| Border Collie x unknown | 9 | 3 | 4 | **3** | **4** |
| Shetland Sheepdog | -- | -- | 4 | **3** | **2** |
| Koolie | -- | -- | 1 | **2** | **< 1** |
| Miniature American Shepherd | -- | 2 | -- | **2** | **< 1** |
| Beauceron | 4 | -- | -- | **3** | **< 1** |
| Beauceron x unknown | 4 | -- | -- | **2** | **< 1** |
| Rough Collie | -- | -- | 1 | -- | **1** |
| Welsh Corgi | -- | 2 | -- | **< 1** | **< 1** |
| **Other breed groups** |  |  |  |  |  |
| Great Dane | -- | 8 | 8 | **< 1** | **7** |
| Catahoula | -- | 2 | 4 | -- | **3** |
| Dachshund | 4 | 2 | 2 | **1** | **2** |
| Chihuahua | 4 | 2 | 1 | **1** | **2** |
| American Cocker Spaniel | -- | -- | 2 | -- | **1** |
| **(b) Sex** |  |  |  |  |  |
| **male** | 39 | 56 | 55 | **51** | **53** |
| **female** | 61 | 44 | 45 | **49** | **47** |
| **(c) Site of acquisition** |  |  |  |  |  |
| **rescue** | 57 | 46 | 59 | **8** | **55** |
| **breeder** | 9 | 5 | 2 | **47** | **4** |
| **private** | < 1 | 16 | 4 | **42** | **7** |
| **no response<sup>a</sup>** | 34 | 33 | 34 | **3** | **34** |
| **(d) Other dog(s) at home** |  |  |  |  |  |
| **yes** | 83 | 70 | 85 | **62** | **80** |

In the list of breeds, the “x” symbols mean “mixed with”. Breed names without “x” symbol are for purebred breeds. “--” means that no dog was concerned.
<sup>a</sup> “no response” was for owners who did not respond to the optional question regarding the site of acquisition of their dog.

All the breeds listed have standard coat colour patterns with only minor areas of white according to kennel clubs. Also, all breeds possibly carry three of the genes (Merle, piebald, Irish spotting) whose mutations are known to cause both pigment deletion in hairs and eyes and congenital hearing impairments (plus vision impairments for Merle) [25; 3–4].

#### Sex

The frequencies of males and females for each group are listed in **Table 3b**. The group with the highest frequency of females (HNVI = 61%) did not significantly differ from that with the lowest frequency (HIVN = 44%; *X^2^* = 1.82, *p* = 1.0). Overall, males and females were likewise balanced in all groups. Thus, differences between genders in health troubles, aggressiveness, interspecific communication and cooperative activities (*e.g*., [26]) may have been equally compensated for in all groups.

#### Site of acquisition

Owners were free to respond to the following optional question: “Site of acquisition of your dog – where does your dog come from?” The response choices were:

- a rescue centre or a foster program
- a professional, registered breeder
- a private individual or a non-registered breeder.

The response frequencies obtained for each group are provided in **Table 3c**. No response was given for 34% of the sensory impaired dogs and 3% of the sensory normal ones. According to the responses for the 358 remaining dogs, sensory impaired dogs mostly came from rescues/fosters, while sensory normal dogs came from either professional, registered breeders or private individuals/non-registered breeders. The numerous missing responses and the low frequency of “professional breeder” responses for sensory impaired dogs can easily be explained. As described in the Introduction, both the sensory impairments and the morphological abnormalities (see section “Morphology” below) of these dogs result from inopportune, sometimes illegal, breeding practices. Consequently, these dogs cannot officially be sold by registered breeders.

#### Presence of congeners at home

Owners were asked to indicate whether they had other dog(s) at home than that concerned by their participation in the survey. This question was asked because many rescue centres and foster programs that propose sensory impaired dogs for adoption recommend the presence of at least one sensory normal dog at the adopter’s home. According to these rescues/fosters, the sensory normal dog is expected to become a “referent” for the sensory impaired dog concerning various aspects of life, such as, for example, spatial exploration and interactions. The responses obtained for each group are presented in **Table 3d**. Significantly more sensory impaired than sensory normal dogs were reported as living with congeners (frequencies = 80% and 62%, respectively; *X^2^* = 17.55, *p* = 0.002). This difference may be explained by the above-mentioned recommendation, provided that many sensory impaired dogs were adopted from rescues/fosters. An additional, related explanation is that the adoption of a sensory impaired dog needs prior experience in dog-human communication and dog training. As a result, sensory impaired dogs are more frequently adopted by persons that already have had, or presently have, dogs. However, we have no hypothesis as to whether the presence of congeners at home could have differently affected the responses for sensory impaired and sensory normal dogs for the various data compared below. This point is therefore not analysed in further detail.

### Determination of sensory impairments

#### Severity of the impairment

As mentioned above, owners were asked to report the sensory status of their dog, at each ear and each eye separately, by choosing one of three possible responses:

- normal
- partially impaired
- totally impaired (deaf/blind).

**Table 4** shows how the responses were used to provide a “severity” score to hearing and vision impairments. Scores 1 and 2 mean that the impairment is unilateral. Scores 3 and 4 mean that both ears/eyes are impaired but at possibly different degrees. Score 5 means that the impairment is both total (*i.e.*, deafness/blindness) and bilateral.

**Table 4.** Score of severity of hearing and vision impairments determined from owners’ responses to the sensory status of their dog at each ear/eye.

| Score of severity | Sensory status at one ear/eye | Sensory status at the other ear/eye |
| --- | --- | --- |
| 1 | normal | partially impaired |
| 2 | normal | totally impaired |
| 3 | partially impaired | partially impaired |
| 4 | partially impaired | totally impaired |
| 5 | totally impaired | totally impaired |

The left panel in **Fig 2a** shows the distributions of severity scores for the two hearing impaired groups. Scores were equally distributed in the two groups (mean scores of severity ± 1 standard deviation for HIVN and HIVI groups = 4.6 ± 0.9 and 4.8 ± 0.7, respectively). Most hearing impaired dogs were reported as being bilaterally deaf (frequencies of score 5 for HIVN and HIVI groups = 81% and 87%, respectively). The right panel in **Fig 2a** shows the distributions of severity scores for the two vision impaired groups. Only 25% of the vision impaired dogs were reported as being bilaterally blind. Thus, most vision impaired dogs had residual, unilateral or bilateral, vision. The two vision impaired groups showed no clear difference in score distributions, in spite of the trend for score 3 to be slightly more frequent for the HIVI group (mean scores of severity ± 1 standard deviation for HNVI and HIVI groups = 3.3 ± 1.5 and 3.4 ± 1.3, respectively).

**Fig 2.**
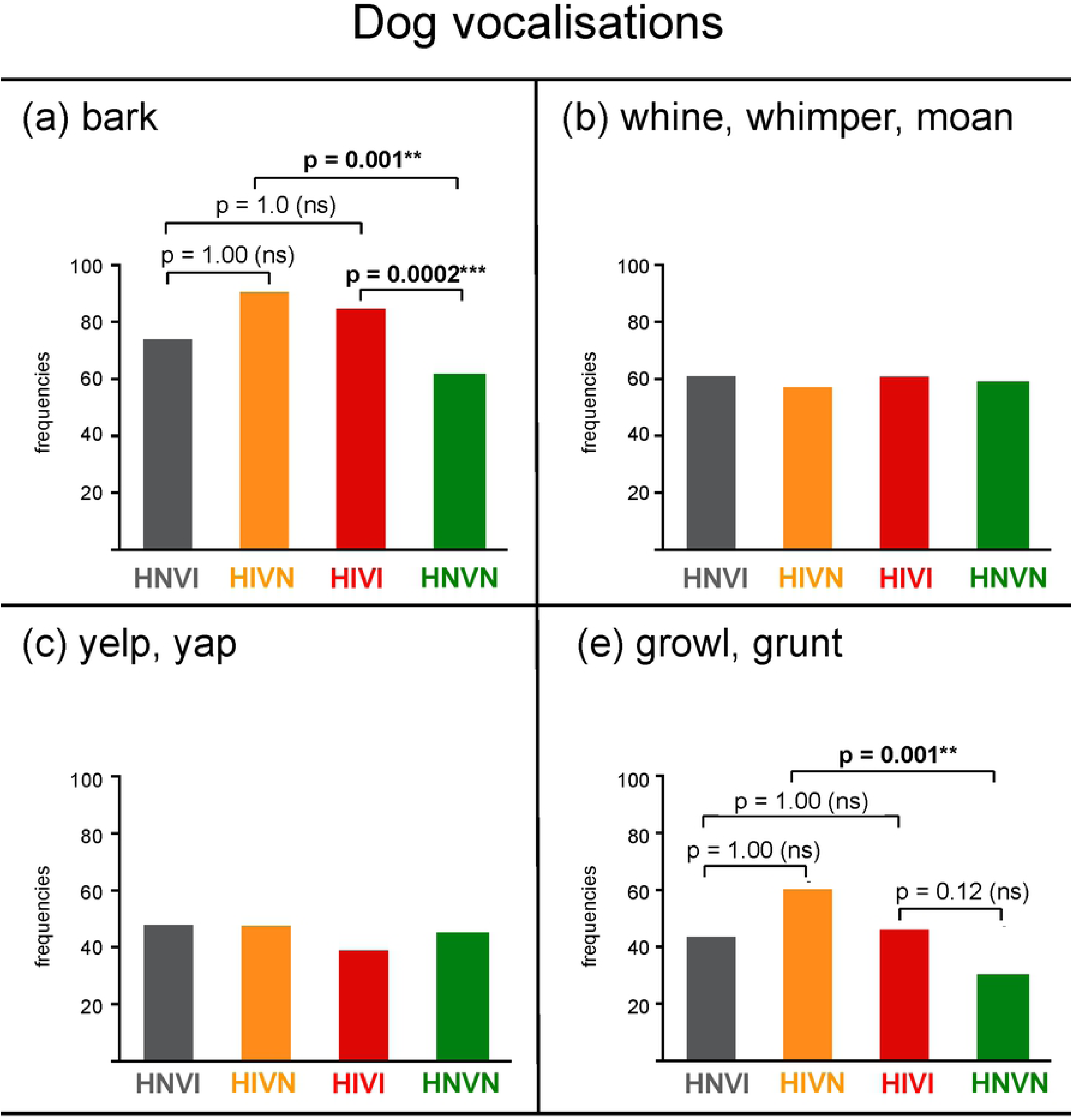
**Frequencies of responses, in percentages, obtained for hearing impaired groups (left panels) and for vision impaired groups (right panels) for the following data: (a) score of severity of the impairment, (b) type of diagnosis test, and (c) operator of the subjective test.** HNVI = hearing normal vision impaired (grey), HIVN = hearing impaired vision normal (orange), HIVI = hearing impaired vision impaired (red).

#### Diagnosis test

Owners who reported sensory impairment(s) in their dog were asked to indicate whether the impairment(s) had been diagnosed using either:

- objective testing (*i.e.*, BAER test in certified clinic for hearing; Canine Eye Registration Foundation – CERF – or equivalent standardised ophthalmic test in certified clinic for vision)
- subjective testing (*i.e.*, someone produced sounds/visual signals and observed the dog’s reaction to these signals).

For vision impairments, an additional response choice was available:

- “diagnosis of vision impairment just based on abnormal eye(s) aspect”. This response was for dogs having severe ophthalmic abnormalities, such as for example no eyeball, which noticeably affect visual function (see pictures of vision impaired dogs in **S1 Fig**; see also section “Ophthalmic abnormalities” below).

The responses obtained for hearing impaired and vision impaired groups are presented in the left and right panels of **Fig 2b**, respectively. Sensory impairments were seldomly diagnosed using objective testing (frequencies of BAER tested dogs for HIVN and HIVI groups = 17% and 8%, respectively; frequencies of CERF-like tested dogs for HNVI and HIVI groups = 17% and 29%, respectively). The finding that hearing impaired dogs were mostly diagnosed using subjective testing can easily be explained by the small number of veterinary clinics that propose BAER testing (see [19] for United States of America and several other countries; see https://www.centrale-canine.fr/lofselect/actualites/la-surdite-comment-la-depister for France). Vision impairments were almost equally diagnosed from CERF-like testing, subjective testing and aspect of the eye(s).

#### Operator of the subjective test

Owners who responded that the sensory impairment(s) of their dog had been diagnosed using subjective testing were asked to indicate who had conducted that subjective test by choosing one the following responses:

- a veterinary
- an employee or a volunteer in a rescue centre/foster
- the owner of the dog (themselves)
- the breeder of the dog.

The responses obtained for hearing impaired and vision impaired groups are presented in the left and right panels of **Fig 2c**, respectively. Subjective testing was performed by a veterinary in 61 to 67% of the cases. Subjective testing was otherwise more frequently performed by owners to evaluate vision than to evaluate hearing. Accordingly, unilateral and/or partial impairments are more easily noticeable when they concern vision. Unilateral hearing impairments mainly affect sound source localisation while having less noticeable effect on sound detection. Unilateral – and/or partial – vision impairments affect the stereoscopic processing of space, objects, human gestures, *etc*, which has a visible impact on both the motion and the posture of the dog. Moreover, subjective testing of monocular vision is much easier to conduct than subjective testing of monaural hearing. A visual source presented on the edge of the visual field can exclusively be processed by the ipsilateral eye. Conversely, a sound reaches the two ears regardless of the spatial position from which it is presented. This explains why BAER testing is currently the only test of unilateral hearing impairments. However, it should be mentioned that most clinical BAER tests use a single sound (*e.g*., a click) presented at either fixed or few different level(s), which does not allow assessing partial hearing impairments at one ear. This could possibly explain in part with it is often stated that congenital hearing impairments in dogs can be unilateral but are always total in the impacted ear [27].

#### Stimuli and conditions of the subjective test

The 171 owners who indicated that the hearing impairment of their dog had been diagnosed using subjective testing were asked to “describe in a few words what the test consisted of”. This open question was asked in order to get insight on the sounds, sites and conditions of the subjective tests that are performed in the numerous dogs that have no access to BAER testing. In total, 109 responses were unexploitable, because either the subjective test had been conducted prior to adoption of the dog by the respondent or the response given was too vague (*e.g*., “my vet made different noise to observe my dog’s reaction”, “my dog has never reacted to any sound”, *etc*).

According to the 62 exploitable responses, most sounds used in subjective testing of hearing were natural sounds. These natural sounds were produced by either clapping/snapping/banging hands or fingers (22 responses), shaking/striking/dropping on the floor a metal object (12 responses), calling/talking to the dog out loud (11 responses), ringing a doorbell or an alarm (5 responses), producing whistles (5 responses), using a clicker (1 response) or a tuning fork (1 response), or turning on a vacuum cleaner (1 response). Seven other respondents indicated that sounds were produced using an automated device, such as a smartphone application or an audiometer, which allowed playing tonal or narrowband “artificial” sounds of different frequencies at several levels. On the subject of the test conditions, 14 respondents indicated that sounds were intentionally produced while the dog was sleeping. These 14 dogs were considered as hearing impaired because the presented sound(s) did not wake them up. Regardless of dog arousal, sounds were produced either very close to the dog’s ear (7 responses) or out of sight from several locations and distances (18 responses). Only one respondent mentioned occlusion of one ear during sound presentation, but without specifying how exactly the ear had been occluded.

### Morphology

The survey was specifically addressed to owners of dogs with no or congenital sensory impairments. As detailed in the Introduction, most congenital impairments in dogs are associated with genetic-related deletion of pigments in hair and irises. Below, we therefore assessed to what extent sensory impairments were associated with discolouration of the coat and irises. Ophthalmic abnormalities, which are consistently reported in dogs with mutation of one pigment deletion gene, namely in homozygous Merles, were also assessed.

#### Excess white coat

Owners were asked to indicate what surface of the dog’s coat was white, on the body and head separately, by choosing one of the following responses:

- less than 50%
- between 50 and 75%
- more than 75%.

Because all dogs belonged to breeds whose standard coat includes only minor areas of white, dogs reported as having 50% or more of white were considered as “excess white”.

The results obtained for each group for the body and head are presented in **Fig 3a** and **Fig 3b**, respectively. Few sensory normal dogs (frequencies for the HNVN group ≈ 10%) but most sensory impaired dogs (frequencies for HNVI, HIVN and HIVI groups ranging from 74% to 97%) had excess white coat. There was a non-significant trend for higher frequencies for the HIVI group than for HNVI and HIVN groups (*X^2^* ≤ 9.75, *p* ≥ 0.07). Pictures of 88 dogs sent by their owners can be seen in **S1 Fig** as illustrative examples of the coat colours most frequently reported for each group.

**Fig 3.**
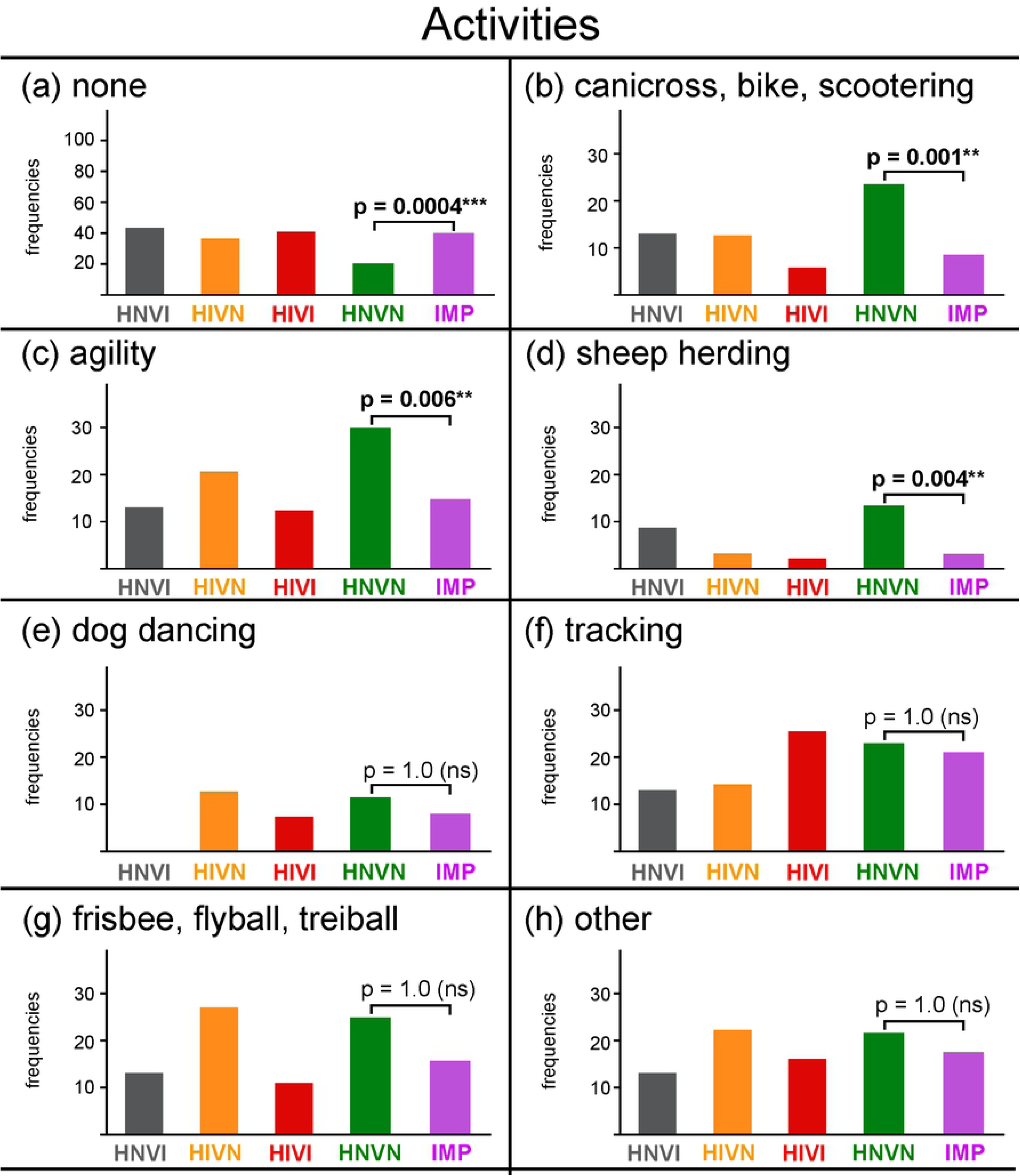
**Frequencies of responses, in percentages, obtained for each group for the following morphological data: (a) excess white coat on body, (b) excess white coat on head, (c) discoloured or indiscernible iris, (d) ophthalmic abnormalities.** HNVI = hearing normal vision impaired (grey), HIVN = hearing impaired vision normal (orange), HIVI = hearing impaired vision impaired (red), HNVN = hearing normal vision normal (green), ns = not significant. Horizontal brackets show two-by-two comparisons assessed using Chi^2^ tests.

#### Iris colour

Owners were asked to indicate whether the colours of the left and right irises of their dog were either:

- normal for the breed standard (*e.g*., brown, green, deep blue)
- discoloured to extreme light blue
- indiscernible (due to absence of eyeball, covering by eyelid or membrane, *etc*).

The frequencies of dogs with discoloured or indiscernible iris were assessed regardless of whether the “discoloured” or “indiscernible” response was selected for one or both eyes.

The results obtained for each group are provided in **Fig 3c**. Few sensory normal dogs (frequency = 12%) but most sensory impaired dogs (frequencies > 80%) had discoloured or indiscernible iris. Frequencies were similar for the two vision impaired groups (91% and 96%, respectively; *X^2^* = 1.20; *p* = 1.0), and were slightly lower for HIVN group (81%; comparison with HIVI group: *X^2^* = 13.16, *p* = 0.01; comparison with HNVI group: *X^2^* = 1.32, *p* = 1.0, *ns*).

#### Ophthalmic abnormalities

Owners were asked to indicate whether their dog had, at the left and right eyes separately, the following ophthalmic abnormalities:

- microphthalmia
- misshapen pupil
- cataract
- absence of eyeball
- other than those mentioned above
- the dog has no, listed or “other”, ophthalmic abnormalities.

Multiple responses were allowed. The list was based on pilot data on 40 excess white dogs collected by the second author and their veterinaries, and was followed by a field for manual report of the type(s) of “other”, non-listed, ophthalmic abnormalities.

The frequencies of dogs having at least one ophthalmic abnormality, regardless of whether this was at one or both eyes, are presented for each group in **Fig 3d**. Ophthalmic abnormalities were seldom for the HNVN group (frequency = 8%) but were extremely frequent for vision impaired groups (frequencies for HNVI and HIVI groups = 83% and 91%, respectively). There was no statistical difference between the two vision impaired groups (*X^2^* = 1.30, *p* = 1.0). Compared to vision impaired groups, the HIVN group showed ophthalmic abnormalities to a significantly smaller frequency (30%; *X^2^* ≥ 18.75, *p* ≤ 0.0005).

**Fig 4** shows the frequencies at which each ophthalmic abnormality was reported for sensory impaired groups. There was no difference between the two vision impaired groups (HNVI *vs.* HIVI: *X^2^* ≤ 1.45, *p* = 1.0). For these two groups, the most frequent abnormality was microphthalmia (frequencies = 70% and 64%, respectively), followed from afar by misshapen pupil, cataract, absence of eyeball, and “other” (frequencies ranging from 12% to 27%). The HIVN group significantly differed from either one or both vision impaired groups in the responses relative to microphthalmia (*X^2^* ≥ 29.39, *p* ≤ 0.0001), cataract (*X^2^* ≥ 8.54, *p* ≤ 0.02) and absence of eyeball (*X^2^* = 9.10, *p* = 0.03), but not in those relative to misshapen pupil (*X^2^* ≤ 3.94, *p* = 1.0) and “other” ophthalmic abnormalities (*X^2^* ≤ 4.45, *p* = 1.0). **Table 5** details the types of “other” ophthalmic abnormalities manually reported by owners.

**Fig 4.**
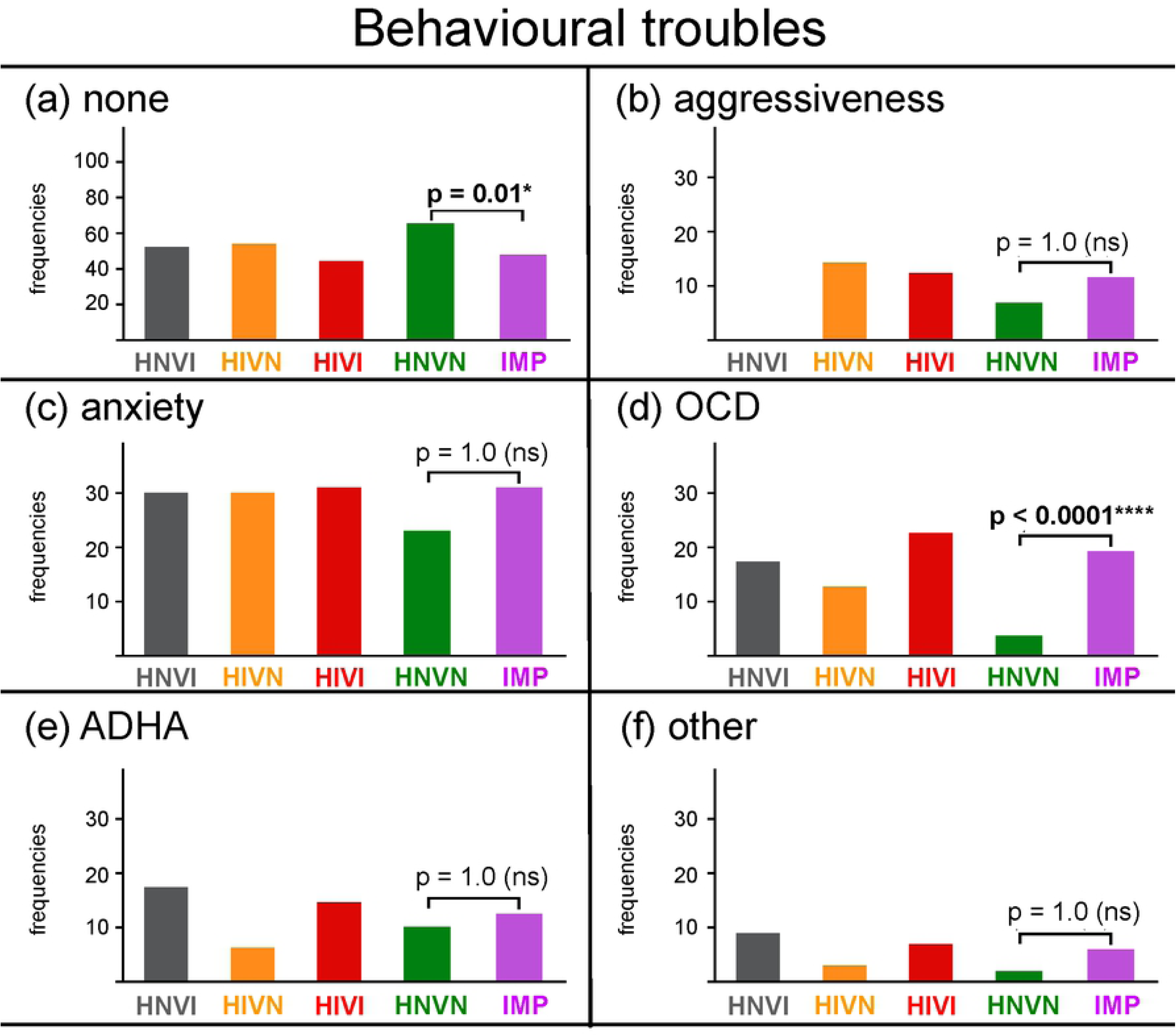
**Frequencies of responses, in percentages, obtained for each sensory impaired group for the following ophthalmic abnormalities: (a) microphthalmia, (b) misshapen pupil, (c) cataract, (d) absence of eyeball, and (e) other.** HNVI = hearing normal vision impaired (grey), HIVN = hearing impaired vision normal (orange), HIVI = hearing impaired vision impaired (red), ns = not significant. Horizontal brackets show the two-by-two comparisons that were statistically assessed using Chi^2^ tests.

**Table 5.** Raw number of dogs from sensory impaired groups obtained for each “other” ophthalmic abnormalities as manually reported by owners.

|  | HNVI | HIVN | HIVI |
| --- | --- | --- | --- |
| coloboma | -- | 1 | 3 |
| corectopia | -- | 1 | 3 |
| detached retina | -- | -- | 3 |
| dropped or fixated pupil | -- | -- | 5 |
| entropion or ectropion | -- | -- | 8 |
| glaucoma | -- | -- | 3 |
| strabismus | -- | 1 | -- |
| unspecified | 3 | 3 | 10 |

Abnormalities are listed in alphabetic order. The “unspecified” response was for owners who responded "other ophthalmic abnormality than those mentioned above" but without specifying the exact type of the abnormality(s). The “--” symbol means that no dog was concerned.

#### Possible genetic cause

All the ophthalmic abnormalities that are depicted in **Fig 4** and **Table 5** are frequently observed in homozygous Merles [7], but are not associated with the two other pigment deletion genes that are possibly present in the breeds under study (piebald, Irish spotting). Among the 160 vision impaired dogs (23 HNVI, 137 HIVI), 131 (82%) had excess white heads, discoloured or indiscernible iris(es), and ophthalmic abnormalities. Thus, we suggest that these 131 dogs were likely double Merles, although few of them have been directly tested as double Merles on the M locus (9) or at least bred from two parents with Merle phenotype according to their owners (38).

### Health troubles

Owners were asked to indicate whether their dog had ever suffered from the following type(s) of health trouble:

- neurological (*e.g*., seizure, epilepsy, *etc*.)
- heart (*e.g*., heart murmur, malformation, *etc*.)
- bones/joints (*e.g*., dysplasia, *etc*.)
- skin
- digestive
- other than those mentioned above
- the dog has never suffered of any, listed or “other”, health troubles.

Multiple responses were allowed. The list was based on both assumptions on the poor health of double Merles (see Introduction) and unpublished data from a survey of 110 presumed double Merle owners conducted by the second author. The list was followed by a field for manual report of “other”, non-listed, troubles. To note, assumptions predict that double Merles also have issues in their reproductive systems [5]. This point has not been investigated in the present study because many excess white dogs with congenital sensory impairments are neutered early so as to avoid at-risk breeding.

**Fig 5a** presents the frequencies of dogs, for each group and for the IMP cohort, with no health trouble reported. These dogs are labelled below as “healthy”. **Fig 5b-5g** present the frequencies at which the different types of heath troubles were reported for the remaining dogs.

**Fig 5.**
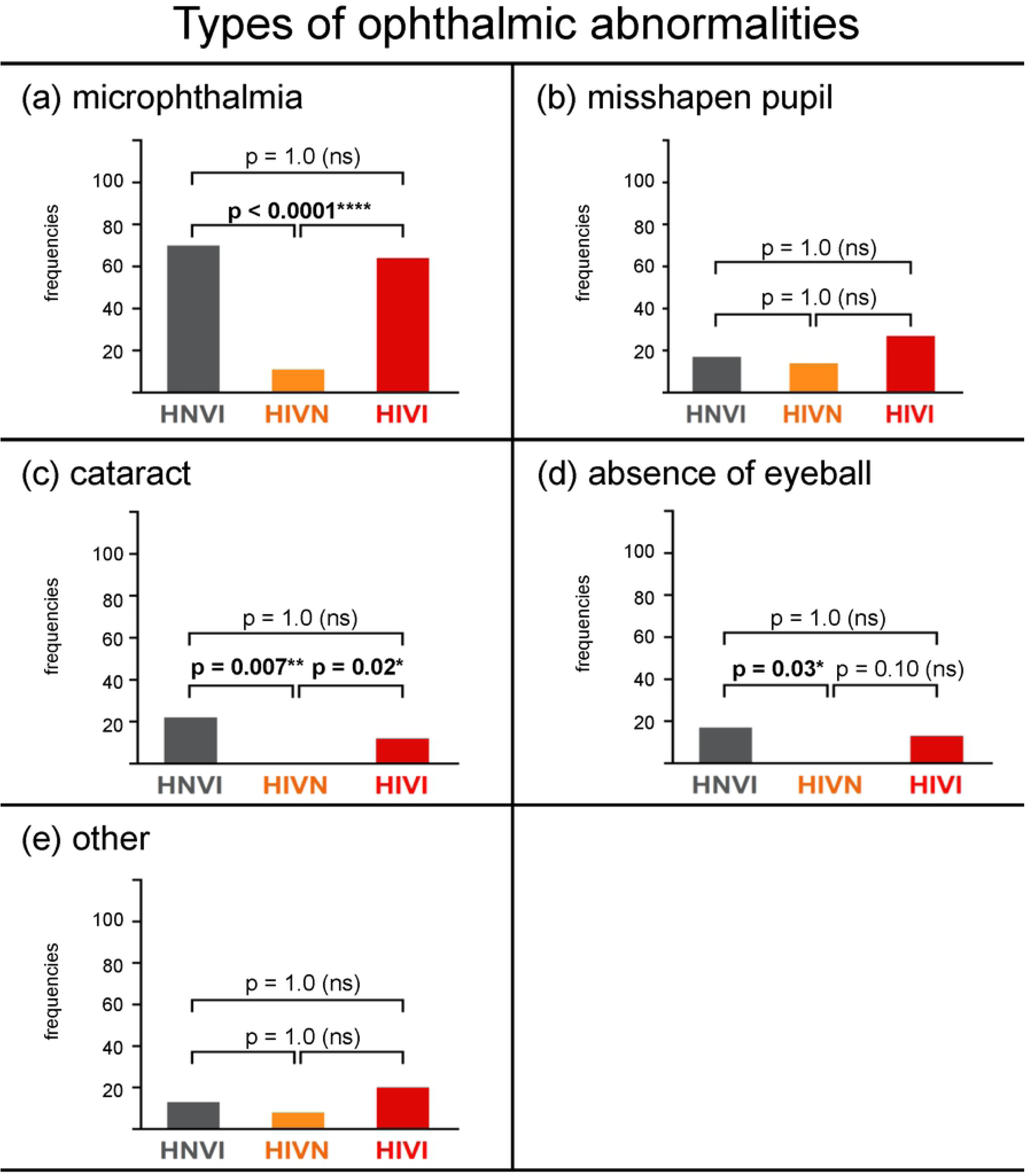
**Frequencies of responses, in percentages, obtained for each group and for the IMP cohort for the following health troubles: (a) none, (b) neurological, (c) heart, (d) bones/joints, (e) skin, (f) digestive, and (g) other.** The ordinate width is larger in panel (a) than in panels (b) to (g). HNVI = hearing normal vision impaired (grey), HIVN = hearing impaired vision normal (orange), HIVI = hearing impaired vision impaired (red), HNVN = hearing normal vision normal (green), IMP = impaired (HNVI, HIVN and HIVI gathered, purple), ns = not significant. Horizontal brackets show comparisons between HNVN and IPM assessed using Chi^2^ tests.

#### Healthy dogs

Seventy-five percent of the sensory normal dogs showed no health trouble according to their owners and were therefore considered as healthy (see Fig 5a). Similar results have been previously reported for comparable breeds in a survey at large scale (*e.g.,* 65% to 75% of “unaffected” dogs within groups of 1,005 Border Collies, 360 Shetland Sheepdogs, and 785 Dachshunds; frequency of unaffected dogs within the group of 71 Australian Shepherds not provided; [23]). Fifty-nine percent of the sensory impaired dogs were healthy, which significantly differs from sensory normal dogs (*X^2^* = 11.86, *p* = 0.02).

#### Neurological troubles

**Fig 5b** shows the frequencies of neurological troubles reported by owners for each group and for the IMP cohort. Overall, neurological troubles were reported for 6.8% of the entire sample. This percentage is substantially higher than those reported in a past survey of about 43,000 dog owners for a large number of diseases of the nervous system (prevalence ≤ 1%, [23]). However, data comparison between the two studies is rendered difficult by several differences. First, Australian Shepherds and Border Collies represented less than 3% of Wiles and colleagues’ sample while representing 82% of the present sample. Second, the survey by Wiles and colleagues exclusively revolved around health, and listed more than 700 specific diseases. In the present study, the section on health troubles was only one of a seven-section questionnaire, and listed only six main categories of health troubles. The two studies had very distinct goals. The study by Wiles and colleagues aimed to quantify the prevalence, across and within breeds, of a variety of specific diseases in the general population of domestic dogs. The small health section in the present study was designed only to assess the veracity of the following assumption: excess white dogs, particularly double Merles, suffer from severe neurological, heart and bones/joints troubles. The present study is therefore not further compared below to that by Wiles and colleagues.

If both the above-mentioned assumption and our suggestion that at least 131 of the sensory impaired dogs in the sample were double Merles were true, then the frequency of neurological troubles reported for sensory impaired dogs (HNVI = 9%, HIVN = 2%, HIVI = 15%, IMP = 11%) should have been higher. However, neurological troubles were significantly less frequently reported for sensory normal dogs (3%) than for sensory impaired ones (*X^2^* = 11.07, *p* = 0.04). Whether – and to what extent – this difference is related to the double Merle genotype, as suggested by the assumption, is undetermined. Reports of neurological troubles indeed mainly concerned vision impaired, possibly double Merle, dogs. Among the 131 dogs presumed above to be double Merles according to their morphological data, 19 (14.5%) had neurological troubles. On the other hand, only two of the 16 sensory impaired dogs that have been tested on the M locus as double Merles showed neurological troubles. Below, we propose two possible, complementary explanations as to why neurological troubles were more frequently reported for sensory impaired dogs than for sensory normal ones.

##### Undiagnosed MDR1-related drug sensitivity

All except three of the 31 dogs for which neurological troubles were reported (25 sensory impaired, 6 sensory normal) were Australian Shepherds, Border Collies or Rough Collies. As mentioned in the Introduction, mutation of the MDR1 gene is frequent in these breeds [14–15]. This mutation prevents the blood-brain barrier from blocking chemical agents at the entrance of the central nervous system. As a result, commonly administered drugs (including antibiotics, anti-diarrheal, parasite control products, pain medications, sedatives and tranquilisers) that rouse no deleterious reaction in dogs with normal MDR1 elicit severe neurological symptoms (*i.e*., seizure, tremors, disorientation) in dogs with mutated MDR1.

Owners were asked to indicate whether their dogs had been tested for the MDR1 gene, and, if so, whether the result indicated either normal or – heterozygous or homozygous – mutated allele(s). The frequency of dogs tested for MDR1 was low in the entire sample (26%), and was significantly lower for sensory impaired than for sensory normal dogs (14% and 38%, respectively; *X^2^* > 50, *p* < 0.00001). The few impaired and normal dogs that have been tested showed statistically similar frequencies of MDR1 mutation, and hence of drug sensitivity (19% and 31%, respectively; *X^2^* = 1.60, *p* = 1.0). The smaller frequency of MDR1 testing for sensory impaired dogs that possibly have ophthalmic abnormalities, sensitivity of the skin and eyes to UVs, *etc*., could be explained by the numerous veterinary exams (sensory impairment diagnosis, ophthalmological tests, *etc*.) and specific equipment (sunglasses, vibrating collar, *etc*.) that their owners and rescue centres already incur. Testing these dogs for the MDR1 mutation could possibly be considered as being of secondary importance.

In summary, we suggest that an undetermined part of the dogs from the entire sample could have been undiagnosed for MDR1-related drug sensitivity. The greater report of neurological troubles for sensory impaired dogs than for sensory normal ones could be partially accounted for by their lower frequency of MDR1 testing, and hence by a greater risk of “missing” their drug sensitivity. Accordingly, **Table 6** presents summary data for the 25 sensory impaired dogs for which neurological troubles were reported. Only three of them have been MDR1 tested.

**Table 6.** Summary data for the 25 sensory impaired dogs for which neurological troubles were reported by their owners.

| Group | Breed | Age (yrs) | Excess white body/head | Abn. of iris(es) colour | Opht. abn. | Objective indication for double Merle | MDR1 tested | OCD diagnosed |
| --- | --- | --- | --- | --- | --- | --- | --- | --- |
| HNVI | AS | 5.9 | Y/Y | Y | N | ? | N | N |
| HNVI | CHI | 2.4 | Y/Y | Y | N | 2 parents are Merle | N | N |
| HIVN | AS | 6.6 | Y/Y | Y | N | ? | N | N |
| HIVN | BC | 1.1 | Y/Y | Y | Y | Dog is M/M | **Y neg** | **Y** |
| HIVI | AS | 6.7 | Y/Y | Y | Y | ? | N | N |
| HIVI | GD | 3.8 | Y/Y | Y | Y | 2 parents are Merle | N | N |
| HIVI | RC | 7.6 | Y/Y | Y | Y | ? | N | N |
| HIVI | AS | 2.0 | Y/Y | Y | Y | ? | N | N |
| HIVI | AS | 9.6 | Y/Y | Y | Y | Dog is M/M | N | N |
| HIVI | BC | 4.1 | Y/Y | Y | Y | ? | N | N |
| HIVI | AS | 6.3 | N/N | Y | Y | ? | N | **Y** |
| HIVI | AS | 9.5 | Y/N | Y | Y | ? | N | **Y** |
| HIVI | AS | 1.6 | Y/Y | Y | N | ? | N | **Y** |
| HIVI | AS | 1.0 | Y/Y | Y | Y | ? | N | N |
| HIVI | BC | 2.5 | Y/Y | Y | Y | ? | N | **Y** |
| HIVI | AS | 3.2 | Y/Y | Y | Y | ? | N | N |
| HIVI | AS x ? | 1.3 | Y/Y | Y | Y | ? | N | N |
| HIVI | AS | 1.0 | Y/Y | Y | Y | ? | **Y neg** | N |
| HIVI | AS | 1.9 | Y/Y | Y | Y | ? | N | **Y** |
| HIVI | AS | 7.7 | Y/Y | Y | Y | ? | N | N |
| HIVI | AS x BC | 1.2 | N/Y | Y | Y | ? | N | N |
| HIVI | CAT | 2.9 | Y/Y | Y | Y | ? | N | N |
| HIVI | AS x BC | 4.0 | Y/Y | Y | Y | 2 parents are Merle | N | N |
| HIVI | BC x ? | 2.2 | Y/Y | Y | Y | ? | N | N |
| HIVI | BC | 3.0 | Y/Y | Y | Y | ? | **Y neg** | N |

Abn = abnormalities/abnormal. Oph = ophthalmic. AS = Australian Shepherd. BC = Border Collie. CHI = Chihuahua. GD = Great Dane. RC = Rough Collie. ? = unknown. Y = yes. N = no. neg = negative (no drug sensitivity related to MDR1 mutation).
Data incompatible with the hypothesis that the dog is double Merle are underlined. Data incompatible with our hypothesis that neurological troubles have been confounded with undiagnosed MDR1 drug sensitivity or undiagnosed OCDs are embolded.

##### Undiagnosed compulsive behaviours

During the last three years, the second author has regularly followed the 40 congenitally sensory impaired dogs that have been rescued by and adopted from her organisation. She has observed behavioural stereotypes, often referred to as obsessive compulsive disorders (OCDs), in many of the dogs followed. For example, several dogs exhibited compulsive spinning, circling, tail chasing, star gazing, excessive barking, *etc* (see examples in first part of **S1 Video**). We suggest that when these types of behaviours are verbally described by owners to veterinaries or dog trainers/behaviourists, they can be considered, in foremost instance, as being possibly symptomatic of a neurological disorder. Accordingly, the first part of **S1 Video** shows the compulsive behaviours of two sensory impaired dogs. Both dogs were foremost considered as exhibiting neurological signs, which has finally been refuted by adequate medical screening. More importantly, both dogs showed no more compulsive behaviour after behavioural adjustments of their owners to their sensory impairments, as instructed by the second author (see second part of **S1 Video**). In other words, we suggest that OCDs are frequent in sensory impaired dogs, so that an undetermined part of their reported neurological troubles could have been confounded with undiagnosed OCDs. Accordingly, it can be seen in **Table 6** that OCDs have been diagnosed (see details in “Behavioural troubles” section below) in only six of the 25 sensory impaired dogs for which neurological troubles were reported.

#### Heart and bones/joints troubles

Excess white, double Merle dogs are assumed to also frequently suffer from cardiac and skeletal troubles [5]. **Fig 5c** and **5d** indicate that reports of heart and bones/joints troubles were statistically similar for sensory impaired and sensory normal cohorts (heart = 5% and 1%, respectively, *X^2^* = 6.17, *p* = 0.45; bones/joints = 4% and 8%, respectively, *X^2^* = 2.85, *p* = 1.0). Results for the vision impaired groups, that include 131 presumed double Merles, are much lower than those expected from the assumption (frequencies of heart and bones/joints troubles ranging from 0 and 7%).

#### Skin, digestive and other troubles

Frequencies of skin, digestive and “other” health troubles reported are presented in **Fig 5e, 5f** and **5g**, respectively. These frequencies, ranging from 4 to 14%, were statistically similar for sensory impaired and sensory normal cohorts (skin: *X^2^* = 8.03, *p* = 0.17; digestive: *X^2^* = 4.15, *p* = 1.0; other health troubles: *X^2^* = 2.89, *p* = 1.0), and confirmed the unpublished results from a survey of 110 owners of excess white and sensory impaired dogs. **Table 7** details the “other” troubles as manually reported by owners. Sensory impaired and sensory normal dogs mainly differed in allergies. However, neither the causes nor the symptoms of these allergies were specified by owners.

**Table 7.**
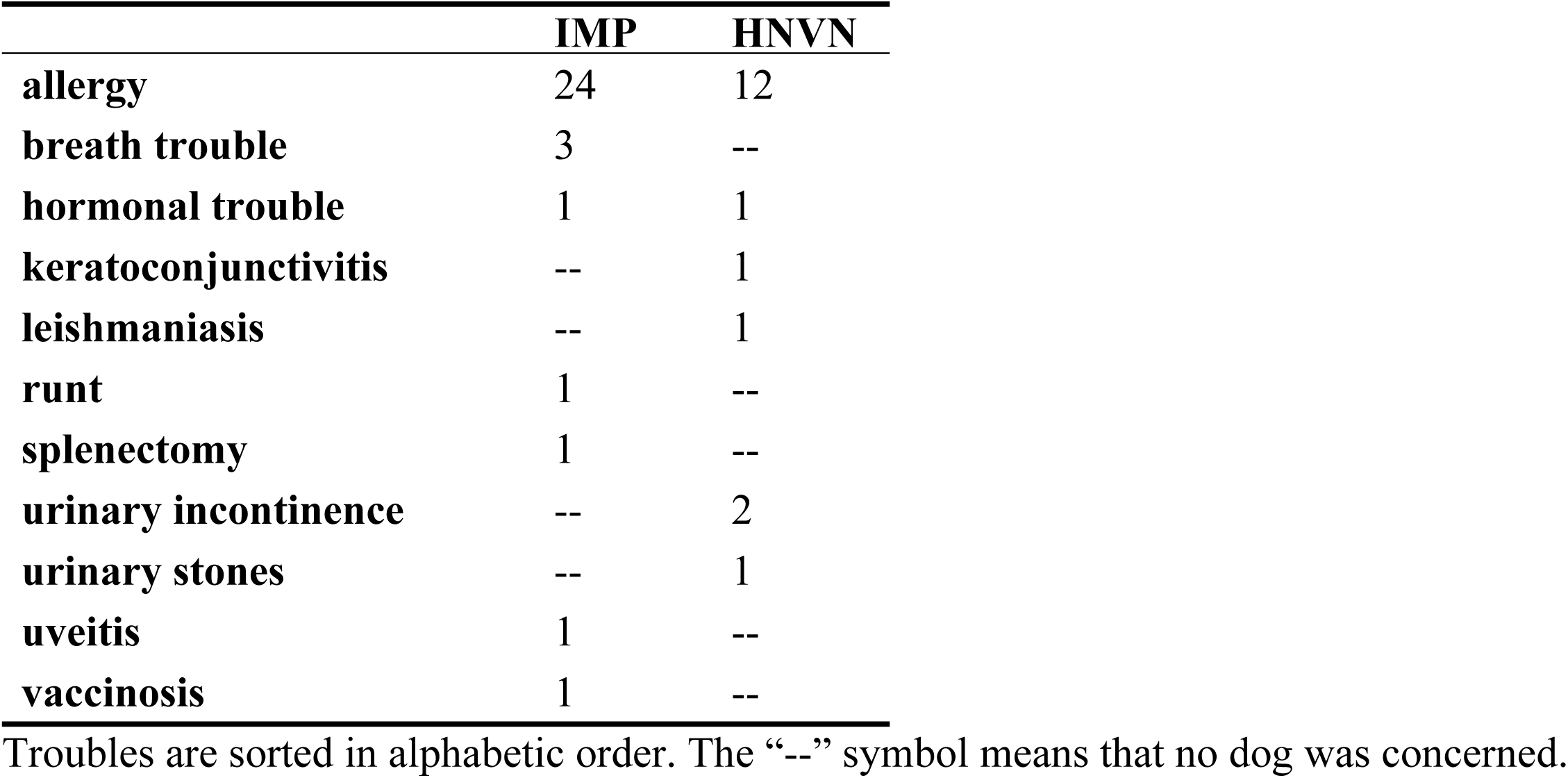
Raw numbers of IMP and HNVN dogs for each “other” (non-listed) health trouble as manually reported by owners.

### Behavioural troubles

Owners were asked to indicate whether their dog had ever suffered from the following behavioural troubles:

- aggressiveness
- anxiety, including separation anxiety
- attention deficit/hyperactivity disorder (ADHD)
- obsessive compulsive disorder (OCD)
- other than those mentioned above
- the dog has never suffered of any, listed or “other”, behavioural troubles.

Multiple responses were allowed. This list was based on:

- the common assumption that deaf and/or blind dogs frequently exhibit aggressiveness and anxiety (see Introduction)
- observations of 40 sensory impaired dogs by the second author during three years (see “Undiagnosed compulsive behaviours” section above)
- informal discussions between the two authors and dog trainers, veterinaries and behaviourists about the behavioural troubles that are frequently observed in Australian Shepherds and Border Collies with insufficient or inadequate activities and interactions.

The list was followed by a field for manual report of “other”, non-listed, troubles. **Fig 6a** presents the frequencies of dogs, for each group and for the IMP cohort, with no behavioural trouble reported. Significantly more sensory normal than sensory impaired dogs had no behavioural troubles (frequencies = 65% and 48%, respectively; *X^2^* = 13.64, *p* = 0.01). **Figs 6b-6f** present the frequencies at which the different behavioural troubles were reported for the remaining dogs.

**Fig 6.**
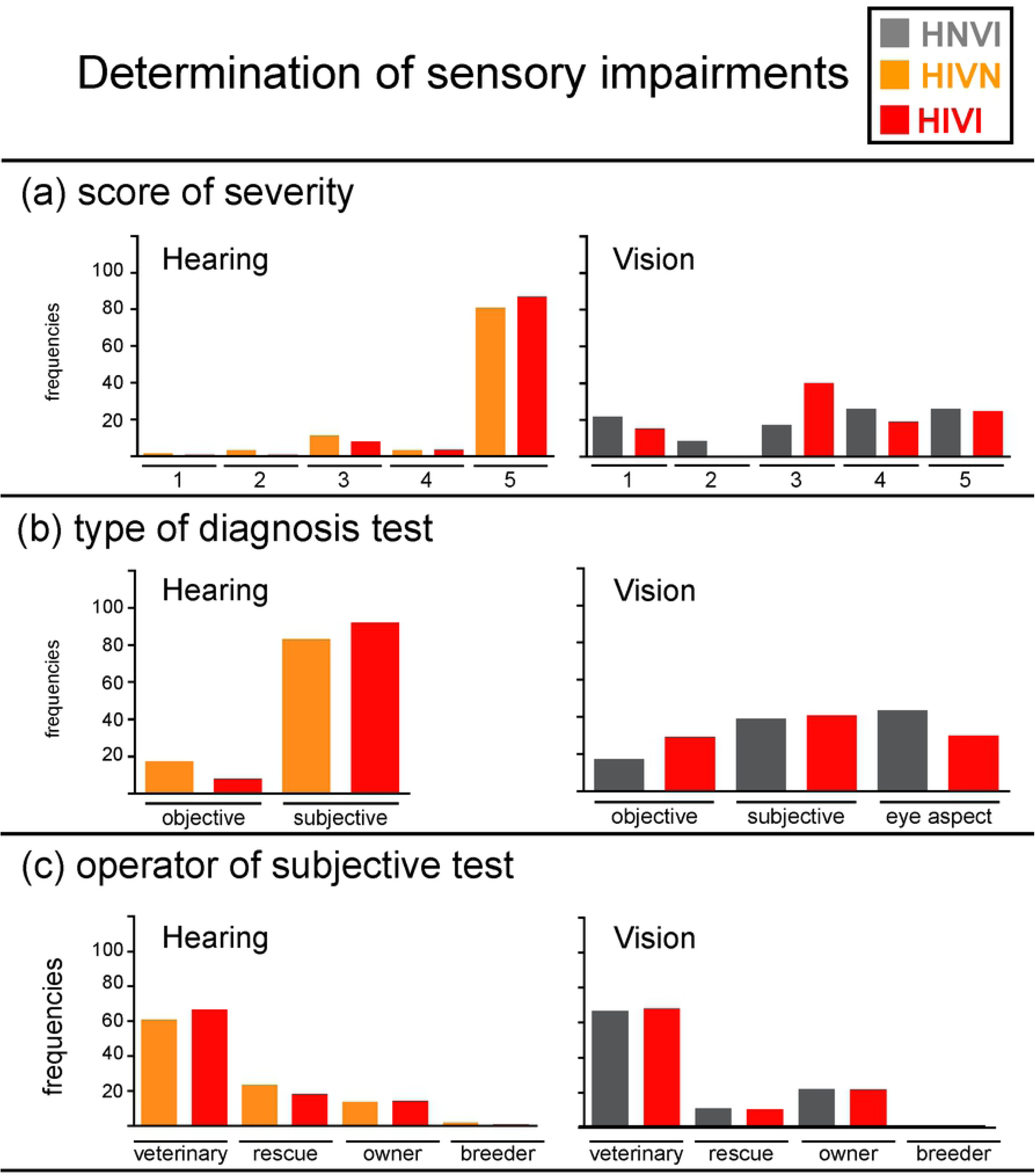
**Frequencies of responses, in percentages, obtained for each group and for the IMP cohort for the following behavioural troubles: (a) none, (b) aggressiveness, (c) anxiety, (d) obsessive-compulsive disorder (OCD), (e) attention-deficit/hyperactivity (ADHD), and (f) other.** The ordinate width is larger in panel (a) than in panels (b) to (f). HNVI = hearing normal vision impaired (grey), HIVN = hearing impaired vision normal (orange), HIVI = hearing impaired vision impaired (red), HNVN = hearing normal vision normal (green), IMP = impaired (HNVI, HIVN and HIVI gathered, purple), ns = not significant. Horizontal brackets show comparisons between HNVN and IPM assessed using Chi^2^ tests.

Aggressiveness was likewise seldom for sensory normal and sensory impaired cohorts (frequencies = 7% and 12%, respectively; *X^2^* = 2.93, *p* = 1.0; see **Fig 6b**), which is opposite to the above-mentioned assumption. There is only one past study that we are aware of that compared behavioural troubles in sensory impaired and sensory normal dogs [22]. As for the present study, the authors conducted an owner survey. However, there are four main differences between the study by Farmer-Dougan and colleagues and the present one. First, the authors used a previously existing questionnaire (*i.e*., Canine Behavioural Assessment and Research Questionnaire, C-BARQ [28]). Second, their respondents had to quantify the severity or frequency of each behavioural trouble listed using 0–4 scales. Third, the authors had no inclusion criteria regarding the type of sensory impairment (*i.e*., congenital or late onset, hereditary or acquired, sensorineural or conductive). Fourth, they investigated a much larger variety of dog breeds (see **Table 3** in [22]). Farmer-Dougan and colleagues found smaller scores of aggressiveness for sensory impaired than for sensory normal dogs, which differs from the present finding of similar frequencies of aggressiveness for both cohorts. However, both studies refute the above-mentioned assumption.

Anxiety was likewise frequent for sensory normal and sensory impaired cohorts (frequencies = 23% and 31%, respectively; *X^2^* = 3.48, *p* = 1.0; see **Fig 6c**). Farmer-Dougan and colleagues found lower anxiety scores for sensory impaired than for sensory normal dogs [22]. Both studies thus refute the above-mentioned assumption. However, Farmer-Dougan and colleagues assessed the behavioural traits that are listed in the C-BARQ, while we have determined our list of behavioural troubles from common assumptions, pilot observations, and informal discussions with professionals. The behavioural data of the two studies are therefore not further compared below. The high prevalence of anxiety in our sensory normal cohort (23%) is similar to that previously reported in various breeds for three items relative to anxiety (*i.e*., separation anxiety, fearfulness and noise sensitivity, see [29]).

Reports of OCDs were seldom for sensory normal dogs (frequency = 4%) but were five times more frequent for sensory impaired dogs (frequency = 19%; *X^2^* = 26.10, *p* < 0.0001, see **Fig 6d**). This finding is in agreement with both past observations by the second author (see examples of OCDs in the first part of **S1 Video**) and our hypothesis that part of the neurological troubles reported for sensory impaired dogs could have been confounded with – impairment-related – undiagnosed OCDs.

ADHDs (frequencies = 10% and 13%, respectively; *X^2^* = 64, *p* = 1.0; see **Fig 6e**) and “other” behavioural troubles (frequencies = 2% and 6%, respectively; *X^2^* = 5.51, *p* = 0.62; see **Fig 6f**) were reported at similar frequencies for sensory normal and sensory impaired cohorts. **Table 8** details the “other” behavioural troubles as manually reported by owners. Excessive barking, as well as certain eating disorders (*e.g*., pica), can be parts of compulsive behaviours.

**Table 8.** Raw numbers of IMP and HNVN dogs for each “other” (non-listed) behavioural trouble as manually reported by owners.

|  | IMP | HNVN |
| --- | --- | --- |
| depression | -- | 1 |
| eating disorder | 7 | 3 |
| excessive barking | 2 | -- |
| sleep disorder | 5 | -- |

Troubles are sorted in alphabetic order. The “--” symbol means that no dog was concerned.

Owners who reported behavioural troubles in their dog were asked to indicate who had “diagnosed” the trouble(s) by choosing one of the following responses:

- a veterinary specialised in behaviour
- a general veterinary
- a dog trainer or a dog behaviourist
- the owner of the dog (themselves).

They were also asked whether drugs had been prescribed for this/these trouble(s). The responses obtained for sensory impaired and sensory normal cohorts are presented in **Table 9**. Almost 60% of the behavioural troubles have been “diagnosed” by owners. Behavioural troubles have been otherwise diagnosed by a general veterinary (27% of sensory impaired dogs) or a dog trainer/behaviourist (33% of sensory normal dogs). Drugs have been prescribed to only 15% of the dogs with behavioural troubles. To note, the behavioural troubles of many sensory normal dogs that have been prescribed drugs have *not* been diagnosed by a veterinary.

**Table 9.** Frequencies of responses, in percentage, obtained for IMP and HNVN dogs concerning behavioural troubles: (a) operator of the diagnosis and (b) drugs prescription.

|  | IMP | HNVN |
| --- | --- | --- |
| **a. Operator of the diagnosis** |  |  |
| veterinary specialised in behaviour | 5 | 1 |
| general veterinary | 27 | 7 |
| dog trainer or behaviourist | 12 | 33 |
| owner | 56 | 58 |
| **b. Drugs prescription** |  |  |
| yes | 16 | 15 |

Frequencies were assessed from the number of dogs for which behavioural troubles were reported by owners.

### Activities

#### Leisure and sport activities

Owners were asked to indicate how frequently their dog was practicing each of the following leisure/sport activities:

- canicross, bikejoring, scootering
- agility
- sheep herding
- dog dancing
- tracking of objects or persons
- frisbee, flyball, treiball

This list included the activities that are mostly practiced worldwide by the breeds under study, and was followed by an open question that allowed reporting all non-listed activities. To provide their responses, owners had to select one of the following response choices:

- several times a day
- once a day
- several times a week
- once a week
- every two weeks
- once a month
- less frequently than once a month
- never.

We considered that the dog was practicing the activity under examination for all responses except “less frequently than once a month” and “never”.

**Fig 7a** presents the frequencies of dogs, for each group and for the sensory impaired groups gathered (IMP), for which no – listed or “other” – activity was reported. **Figs 7b-7h** present the response frequencies obtained for each activity. **Table 10** details the “other” activities as manually reported by owners.

**Fig 7.**
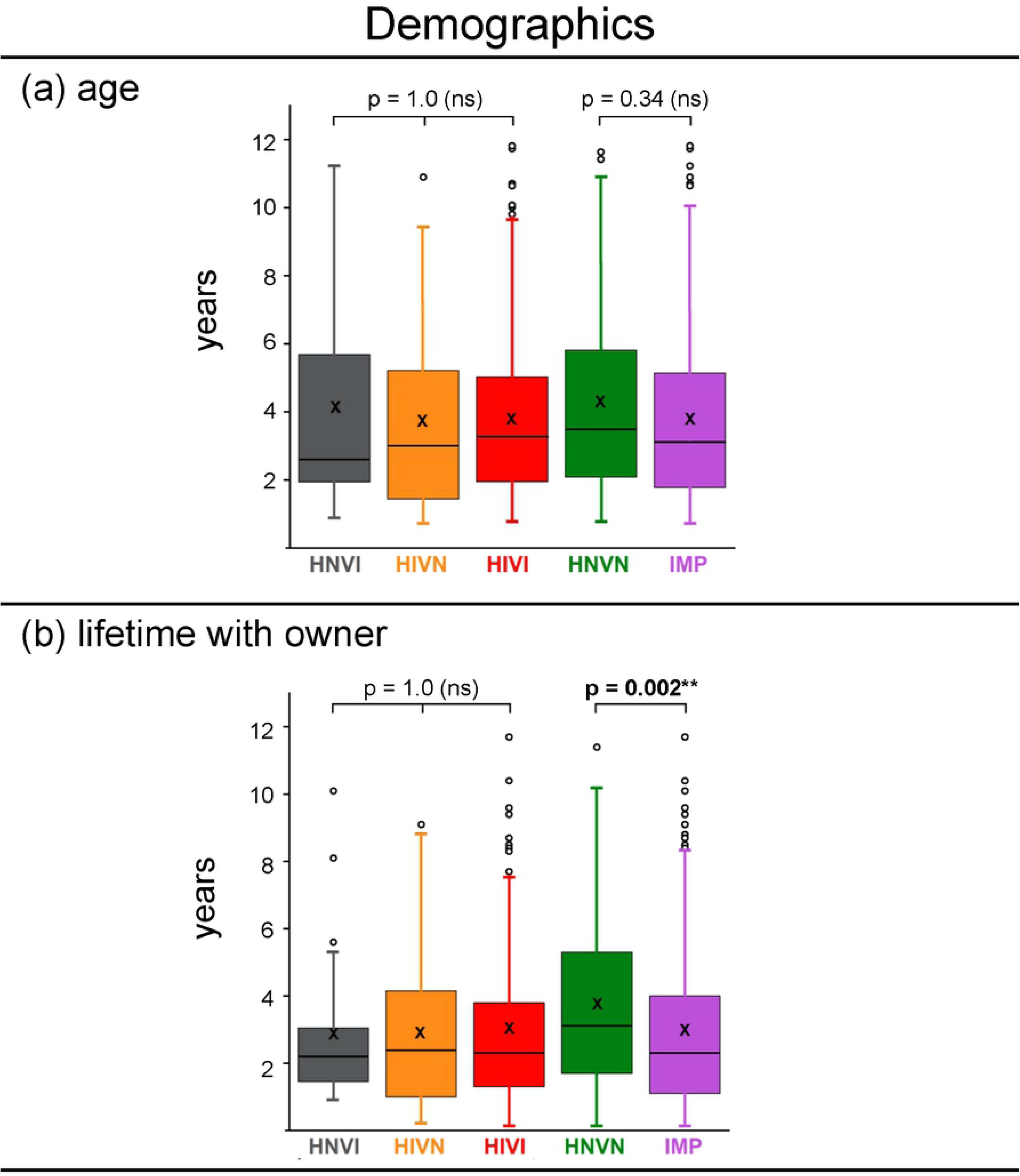
**Frequencies of responses, in percentages, obtained for each group and for the IMP cohort for the following leisure/sport activities: (a) none, (b) canicross/bikejoring/scootering, (c) agility, (d) sheep herding, (e) dog dancing, (f) tracking of objects or persons, (g) Frisbee/flyball/treiball, and (h) other.** The ordinate width is larger in panel (a) than in panels (b) to (h). HNVI = hearing normal vision impaired (grey), HIVN = hearing impaired vision normal (orange), HIVI = hearing impaired vision impaired (red), HNVN = hearing normal vision normal (green), IMP = impaired (HNVI, HIVN and HIVI gathered, purple), ns = not significant. Horizontal brackets show comparisons between HNVN and IPM assessed using Chi^2^ tests.

**Table 10.** Raw numbers of IMP and HNVN dogs for each “other” (non-listed) activity as manually reported by owners.

|  | IMP | HNVN |
| --- | --- | --- |
| **Barn hunt** | 2 | 1 |
| **Cani-roller** | -- | 1 |
| **Cani-walk** | -- | 1 |
| **Dog diving** | 1 | -- |
| **Hiking** | 8 | -- |
| **Hoopers** | -- | 3 |
| **Jumping** | 1 | -- |
| **Kayak** | -- | 1 |
| **Lure course** | 2 | 1 |
| **Nosework\*** | 5 | 1 |
| **Obedience** | 2 | 13 |
| **Paddle** | -- | 4 |
| **Paragliding** | -- | 1 |
| **Parkour** | 1 | 1 |
| **Rally-O** | 3 | 2 |
| **Retrieving\*** | 1 | 2 |
| **Ring** | -- | -- |
| **Seek\*** | 2 | -- |
| **Skijoring** | 1 | -- |
| **Sled** | 1 | -- |
| **Swimming** | 6 | 8 |
| **Trail running** | 4 | 3 |
| **Tricks** | 9 | 2 |

Activities are sorted in alphabetic order. Activities followed by an asterisk strongly rely on olfactory capacities. The “--” symbol means that no dog was concerned.

Twenty percent of the sensory normal dogs, against 40% of the sensory impaired dogs, were involved in absolutely no canine activity according to their owners (*X^2^* = 20.10, *p* = 0.0004). This large difference can easily be explained by both the assumption that sensory impaired dogs are poorly capable of practicing activities and the fact that many official competitions in the countries under study have long been inaccessible to dogs that are sensory impaired and/or unregistered in kennel clubs. Accordingly, the following three activities were significantly more frequently practiced by sensory normal dogs than by sensory impaired ones: canicross/bikejoring/scootering (frequencies = 24% and 9%, respectively; *X^2^* = 18.45, *p* = 0.001), agility (frequencies = 30% and 15%, respectively; *X^2^* = 14.59, *p* = 0.006) and sheep herding (frequencies = 13% and 3%, respectively; *X^2^* = 15.31, *p* = 0.004). However, the following four activities were practiced at statistically comparable frequencies by sensory normal and sensory impaired dogs: dog dancing (frequencies = 12% and 8%, respectively; *X^2^* = 1.48, *p* = 1.0), tracking (frequencies = 23% and 21%, respectively; *X^2^* = 0.25, *p* = 1.0), frisbee/flyball/treiball (frequencies = 25% and 16%, respectively; *X^2^* = 5.76, *p* = 0.56) and “other” (frequencies = 22% and 17%, respectively; *X^2^* = 1.22, *p* = 1.0). It is noteworthy that tracking, the activity that sensory impaired dogs practiced the most, as well as three other activities listed in **Table 10** (*i.e*., nosework, retrieving, seek), essentially rely on olfactory capacities. It is also noteworthy that 58% of the sensory impaired dogs that practiced no activity, against 36% of the sensory normal dogs that practiced no activity, exhibited behavioural troubles.

For each above-listed activity, owners were also asked to indicate the dog’s level in that activity by choosing one of the following responses:

- not concerned, because response “never” or “less frequently than once a month” given above
- just for fun, at home or during walks
- beginner in a club
- intermediate in a club
- experienced in a club
- competition/championship.

**Table 11** shows the frequencies of “high level” responses (*i.e*., responses “experienced” and “competition/championship” gathered) obtained for sensory impaired and sensory normal cohorts. No general pattern emerges from these data. Compared to those for sensory normal dogs, frequencies of high level responses for sensory impaired dogs were lower for agility, dog dancing and tracking, were conversely higher for frisbee, and were similar for canicross and sheep herding.

**Table 11.** Frequencies, in percentages, of HNVN and IMP dogs for which the level in the activity practiced was reported as being either “experienced” or “competition/championship”.

|  | HNVN | IMP |
| --- | --- | --- |
| **Canicross** | 6 | 5 |
| **Agility** | 30 | 18 |
| **Sheep herding** | 14 | 14 |
| **Dog dancing** | 12 | 6 |
| **Tracking** | 10 | 6 |
| **Frisbee/flyball/treibball** | 8 | 12 |

Frequencies were assessed from the number of dogs from each group that practice the activity at a minimum frequency of once a month.

#### Assistance and therapy activities

Owners were asked to indicate whether their dog was involved in assistance/therapy activities with:

- elderly persons or groups
- a blind person
- a diabetic or epileptic person, with the role of detecting crises and alerting.

Responses indicated that no dog was engaged in activities with blind persons. Eight percent of the sensory impaired dogs, against 4% of the sensory normal dogs, were involved in therapy/assistance activities with elderly persons or groups. Only two dogs in the entire sample, both being hearing and vision impaired (HIVI), were involved with diabetic/epileptic persons. Accordingly, in the study by Farmer and colleagues, about 3% of 183 hearing and/or vision impaired dogs had therapy or working – rather than family pet – roles at home [22]. It is noteworthy that the ability of assistance dogs to detect epileptic and diabetic crises is based on their ability to perceive small variations in the chemical signals produced by the human’s body, and thus on their olfactory capacities.

### Interspecific communication

Dogs with congenital hearing and/or vision impairments are often believed to have poorer abilities to communicate with congeners and humans. Another belief is that because they had no possibility to benefit from auditory-based vocal learning during early ontogenesis, congenitally deaf dogs are less “talkative” than sensory normal ones. The present study focused on communication with humans, provided the various social and medical roles that dogs are acknowledged to play in working activities with humans. We investigated two aspects of dog-human communication: vocalisations addressed by the dog to the owner during interactions, and communication/training signs addressed by the owner to the dog.

#### Dog vocalisations

Owners were asked to answer to the following question: “Is your dog talkative with you? In other words, which of the following vocalisations does your dog frequently produce in order to communicate with you?

- barks
- whines, whimpers, moans
- yelps, yaps
- growls, grunts
- other than those mentioned above
- “your dog never produces any vocalisation during interactions with you”.”

Multiple responses were allowed. This list of canine vocalisations was based on literature (see review in chapter on communication in [21]), and was followed by a field for manual report of “other”, non-listed, vocalisations.

The responses “no vocalisation” (9 and 4%, respectively, of sensory normal and sensory impaired cohorts) and “other” (4% of both sensory normal and sensory impaired cohorts) were infrequently chosen. **Fig 8** shows the response frequencies obtained for each vocalisation listed. Whines/whimpers/moans (frequencies = 57 to 61%) and yelps/yaps (frequencies = 39 to 48%) were reported at similar frequencies for all groups. However, barks (frequencies for HNVI, HIVN, HIVI and HNVN groups = 74, 90, 85 and 62%, respectively) and growls/grunts (frequencies = 43, 60, 46 and 30%, respectively) were significantly more frequently reported for hearing impaired dogs than for sensory normal ones (*X^2^* ≥ 18.58, *p* ≤ 0.001). One exception to this is noted for the non-significant difference between HIVI and HNVN groups in growls/grunts (*X^2^* = 8.79 *p* = 0.12). The two hearing impaired groups did not statistically differ from the HNVI group (*X^2^* ≤ 3.85, *p* = 1.0). Thus, the present data do not confirm the assumption that congenitally deaf dogs are less talkative than sensory normal or vision impaired dogs.

**Fig 8.**
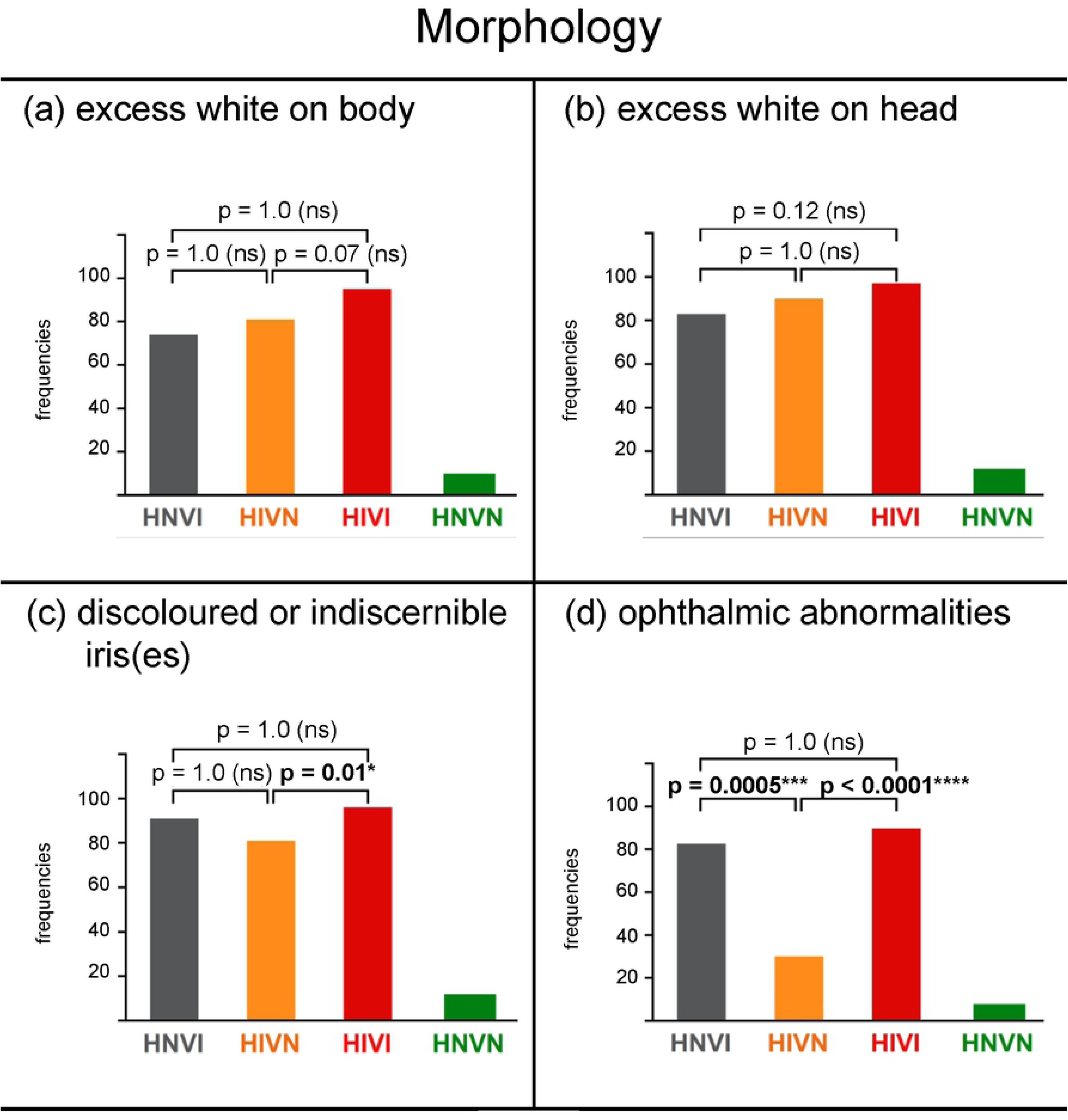
**Frequencies of responses, in percentages, obtained for each group for the following dog vocalisations: (a) barks, (b) whines, whimpers, moans, (c) yelps, yaps, and (d) growls, grunts.** HNVI = hearing normal vision impaired (grey), HIVN = hearing impaired vision normal (orange), HIVI = hearing impaired vision impaired (red), HNVN = hearing normal vision normal (green), ns = not significant. Brackets show the two-by-two comparisons that were assessed using Chi^2^ tests following visual inspection of the data.

#### Human signs

There are four main types of signs that humans can use to communicate with, and train, dogs:

- Gesture, which includes arm, hand, finger or object position and movement, as well as hand sign language
- Sounds, which includes natural and artificial sounds, such as voice, whistle, clicker, *etc*
- Touch, which includes direct touch of the dog’s body with the hand or a stick, remote-controlled vibrating collars, *etc*
- Odours, which includes all odour sources that are manipulated by owners for interactions with their dogs, such as smelling boxes, food pieces, clothes, *etc*.

Owners were asked to indicate which sign(s) they used with their dogs by choosing one response within a long list of unique signs, and combinations of two, three and four above-listed signs (see **S2 Fig**). **Fig 9** shows the responses obtained for each group. For HNVN dogs, the most frequent response was for the “classical” combination of gesture and sounds (frequency = 62%), followed from afar by the combination of all four signs (frequency = 32%). For HNVI dogs, the most frequent response was for sounds only (frequency = 48%), followed by the combination of all four signs (frequency = 26%). For HIVN dogs, the most frequent response was for gesture only (frequency = 63%), followed from afar by the combination of gesture and touch (frequency = 22%). For HIVI dogs, responses were distributed between the touch and odour combination (frequency = 38%), gesture only (23%), touch only (13%), the gesture and touch combination (12%), and the combination of all four signs (9%). In summary, “preferred” signs clearly emerged for dogs with either no or one sensory impairment, but not for dogs with both hearing and vision impairments. Gesture, either alone or in combination with another sign, was almost never used by owners of HNVI dogs, in spite of the large number of dogs with residual vision (see right panel in **Fig 2a**). Odours were almost exclusively used by owners of HIVI dogs, in combination with touch. Thus, odours were almost never used by owners of HNVI and HIVN dogs as communication/training signals, in spite of the different olfaction-based activities in which many of these sensory impaired dogs were involved.

**Fig 9.**
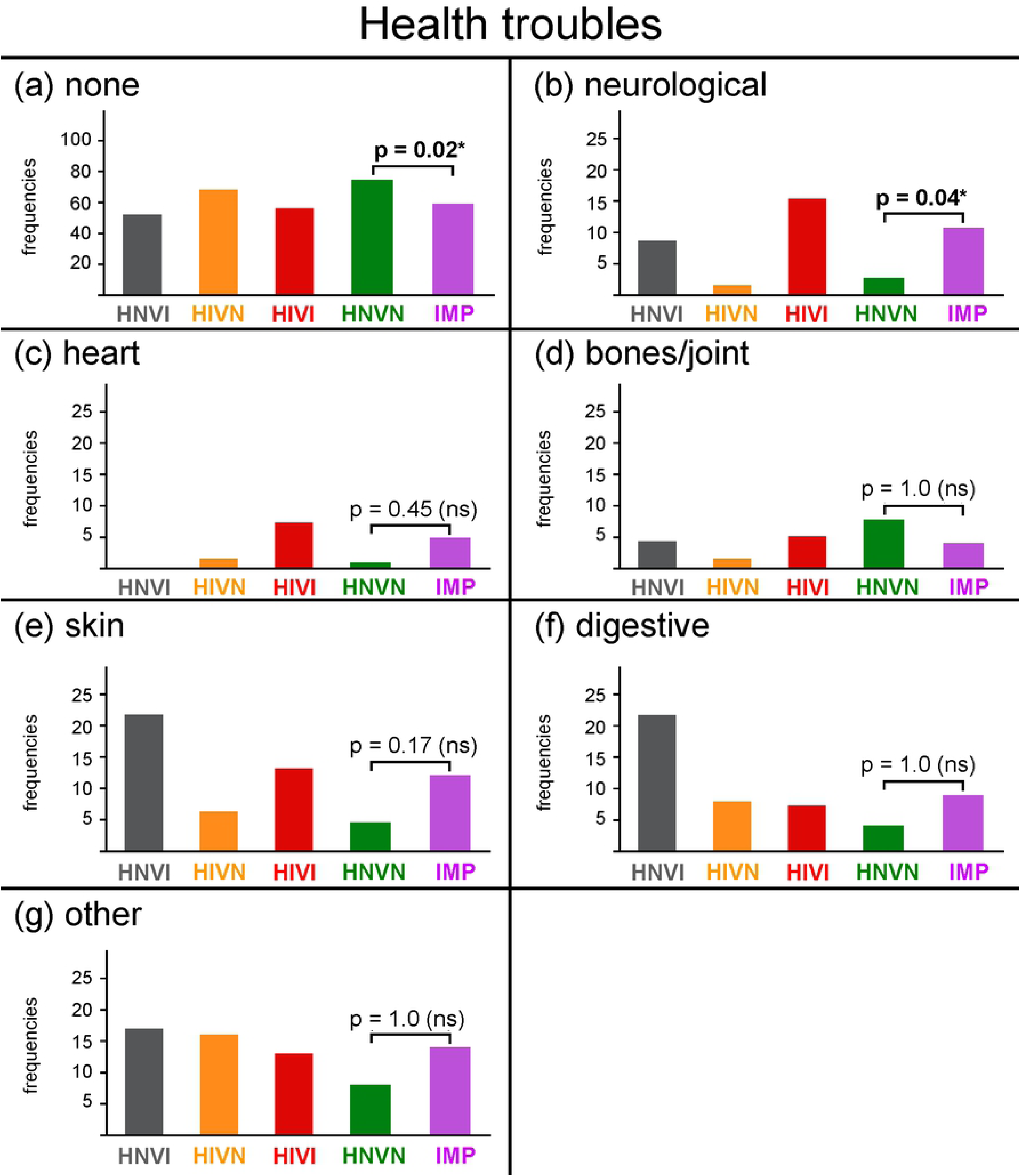
**For each group (panels), frequencies at which different unique signs (left) and combinations of signs (right) were used by owners to communicate with their dogs.** HNVN = hearing normal vision normal (green), HNVI = hearing normal vision impaired (grey), HIVN = hearing impaired vision normal (orange), HIVI = hearing impaired vision impaired (red).

## Summary and conclusions

In this study, we addressed online an international questionnaire to owners of dogs with either no or congenital hearing and/or vision impairments. The main goal of the study was to gain insight on the veracity of various popular assumptions concerning congenitally sensory impaired dogs, that often have dramatic consequences on the future of these dogs (*i.e*., early euthanasia, placement in rescue centres or foster programs with strict adoption criteria, or lack of activities and interactions after adoption). In addition, we aimed to examine both the tools used for determination, and the pigment deletion genetic causes, of the sensory impairments.

### Demographics

As expected, the present study compared two cohorts of congenitally sensory impaired and sensory normal dogs, respectively, that were well matched in size, age, lifetime with owner, breed, and sex. All breeds were possibly concerned by three genes, namely Merle, piebald and Irish spotting, whose mutations are known to produce pigment deletion in hairs and irises and congenital hearing impairments (plus, for Merle, ophthalmic abnormalities associated with vision impairments). The main demographic difference found between the two cohorts was about the site of acquisition of the dog, which was explained by the fact that the births of congenitally sensory impaired dogs often result from irregular breeding.

### Determination of sensory impairments

Hearing impairments were often reported as being both total and bilateral, but were infrequently diagnosed using objective, BAER testing. Veterinary clinics that propose BAER testing are not numerous. Most BAER tests cannot evaluate partial hearing impairments at one ear. Owner responses indicated that subjective testing of hearing was not always performed by a veterinary, and, regardless of the operator, never fulfilled the following three criteria:

- monaural testing with total occlusion of one ear
- presentation of different sounds with various spectral characteristics and levels
- absence of non-auditory – visual and nearfield/floor vibration – cues.

Therefore, we suggest that the capacity of subjective tests to accurately distinguish unilateral from bilateral and partial from total hearing impairments, and hence the reliability of owner reports of these hearing impairments, are low. Vision impairments were almost equally diagnosed using objective (CERF-like) testing, subjective testing, and abnormal aspect of the eye. Most vision impaired dogs in the sample had residual vision.

### Morphology and possibly responsible genes

Coat with excessive white was almost systematically reported for sensory impaired dogs, although sensory impaired and sensory normal cohorts both belonged to breeds in which the normal, standard coat colour pattern is not predominantly white. In addition to this discoloration of the coat, sensory impaired dogs frequently had discoloured or indiscernible iris(es). Both findings are compatible with the hypothesis that the sensory impairments reported had a pigment deletion genetic basis. Normally sighted but hearing impaired dogs showed much fewer ophthalmic abnormalities than vision impaired dogs while showing equally frequent discoloration of the coat and iris(es). Thus, vision impairments in the present sample were likely related to ophthalmic abnormalities. We explained in the Introduction that the mutations of four canine genes, namely Merle, piebald, Irish spotting and KIT, are known to produce pigment deletion in hairs and irises, as well as hearing impairments as a result of a lack of pigments in the stria vascularis of the inner ear. Merle, piebald and Irish spotting are all possibly present in the breeds examined, while KIT only occurs in German Shepherds and is therefore not considered here. Among the three remaining genes, only Merle is additionally associated with ophthalmic abnormalities and concomitant vision impairments. Although few sensory impaired dogs were tested on the M locus as homozygous, double Merles, we suggested that at least 85% of the vision impaired dogs were likely double Merles. If this were true, then the lower number of HNVI dogs compared to that of HIVI dogs could indicate that Merle-related hearing issues are more frequent than Merle-related vision issues. Alternatively, many congenital ophthalmic abnormalities in double Merles are susceptible to worsen over age, and hence to result in a growing, or even late onset, impact on vision. This could not solely explain why so few dogs in the sample only had vision impairments, but also why total blindness was seldomly reported. Further research is needed to quantify the exact prevalence of excess white coat, ophthalmic abnormalities, hearing impairments and vision impairments within a large sample of dogs of various breeds and ages that have all been tested as homozygous for Merle and non-carriers for piebald (as no genetic test is yet available for Irish spotting, and KIT is exclusively present in German Shepherds).

### Health troubles

Significant differences between sensory impaired and sensory normal dogs in health troubles were found for neurological troubles only. Based on morphological data, we have suggested above that (i) most sensory impairments under study were related to pigment deletion gene(s), and (ii) most vision impaired dogs – with either impaired or normal hearing – were likely double Merles. In that event, health troubles were less frequently reported for vision impaired dogs than expected from the common assumption that double Merles suffer from neurological, heart and bones/joints troubles. We propose that the greater report of neurological troubles for sensory impaired dogs than for sensory normal ones may be partially accounted for by their greater lack of diagnosis of both breed-related drug sensitivity and impairment-related compulsive behaviours. Overall, the present data do not confirm the above-mentioned assumption. As explained in the Introduction, this assumption essentially results from multiple citations of a single study [5] that just contained a short statement supported by the citations of few and outdated studies [16–18]. Further research is needed to either refute or confirm assumptions of the poor health of double Merles. The best manner to proceed would be to assess a detailed list of various diseases in a large number of dogs of various breeds and ages that have all been tested as homozygous for Merle, non-carriers for piebald and normal for MDR1, as well as diagnosed for compulsive behaviours.

### Behavioural troubles

Aggressiveness was never reported for HNVI dogs, but was reported at similarly low frequencies for HIVN, HIVI and HNVN dogs. Anxiety was high in all groups. These two findings refute the common assumption that deaf and/or blind dogs exhibit greater aggressiveness and anxiety as a result of the greater frustration caused by their sensory impairments. Prevalence of anxiety in the entire sample is in agreement with past studies. The only difference found between sensory impaired and sensory normal dogs in behavioural troubles was for OCDs, which were considerably more frequent for sensory impaired dogs. This finding is compatible with (i) pilot observations by the second author of frequent OCDs in 40 sensory impaired dogs, and (ii) our hypothesis that undiagnosed OCDs in sensory impaired dogs could have been foremost considered as neurological signs. Responses relative to the diagnosis and medication of behavioural troubles showed that the different behavioural troubles reported by owners were not frequently considered to be severe enough to require professional consultation and chemical treatment, and were otherwise treated using a non-chemical, possibly behavioural, approach.

### Activities

It is generally assumed that sensory impaired dogs cannot be safely and efficaciously engaged in any activity. In the present study, a total lack of activity was twice more frequently reported for sensory impaired dogs than for sensory normal ones. This finding likely reflects the deleterious impact that the general assumption has on the quality of life of sensory impaired dogs. However, the present results indicated that specific leisure activities were practiced at either smaller or equivalent frequencies/levels by the two cohorts. Assistance/therapy activities were even more frequently practiced by sensory impaired dogs. In other words, contrary to the general belief, sensory impaired dogs may be as capable as sensory normal ones of both practicing and achieving good levels of competence in the activities in which their owners engage them. Accordingly, an increasing number of competitions, non-competitive activities and certifications are rendered open to deaf dogs in United States of America (see list in [19]). These positive outcomes may hopefully encourage more owners to engage their sensory impaired dogs in canine activities, which would ultimately reduce the difference between sensory impaired and sensory normal “inactive” dogs. It is noteworthy that most dogs in the entire sample belonged to herding breeds, for which the need for regular physical and mental activities to prevent behavioural troubles related to frustration or boredom (*e.g*., anxiety, ADHD, OCD) has largely been proven. Greater involvement of sensory impaired dogs in activities may therefore have the beneficial effect of reducing their behavioural troubles. Accordingly, recent studies have demonstrated the inverse relationship between engagement in activities and behavioural troubles in sensory normal dogs [30–31].

Moreover, the results indicated that sensory impaired dogs can actually be engaged in both leisure/sport and therapy/assistance cooperative activities that rely on olfactory capacities. There are numerous studies of olfactory capacities in dogs, due to the important social and medical roles that these capacities can play for humans (*e.g*., rescue of missing or enshrouded persons, detection of cancer cells, explosives and toxic fumes, *etc*., see review in [21]). However, there is no data on olfactory capacities in dogs with congenital hearing and/or vision impairments. Brain plasticity during early ontogenesis could possibly have resulted in overdeveloping their olfactory capacities. We suggest that not solely sensory impaired dogs should not be excluded from, but may also exhibit super normal capabilities in, olfaction-based cooperative activities with humans. This that not mean, of course, that we encourage at-risk breeding or births of congenitally sensory impaired dogs. Instead, we expect that present and future research will ultimately reduce the numbers of early euthanasia, placements in rescue centres, and adoptions in overprotective environments, of the numerous sensory impaired puppies that are still born despite the recent developments of knowledge on canine genetics.

### Interspecific communication

The results indicated a trend for hearing and vision impaired dogs to produce more barks and growls/grunts during interactions with their owners than sensory normal dogs. This finding is opposite to the assumptions that congenitally deaf dogs are less “talkative” and that sensory impaired dogs are less capable of communicating with their owners. However, the present study is, to our knowledge, the first attempt to investigate vocalisations in sensory impaired dogs. We cannot determine whether respondents to our survey actually understood the vocalisation terminology used in the questionnaire, whether the vocalisations reported actually had interspecific communication functions, and what emotional valence and arousal had the different vocalisations reported. Also, whether greater barking for sensory impaired dogs is related to compulsive behavioural troubles is undetermined.

Responses concerning human signs to dogs showed that owners are capable of adapting their behaviours to the sensory status of their dogs so as to efficiently communicate with, and train, them. Similar conclusions have previously been drawn from the results of an owner survey [22]. The common assumption that sensory impaired dogs cannot be trained is therefore refuted. To note, for sensory impaired dogs, olfaction was more frequently used in canine activities than in owner communication/training signs.

## Acknowledgments

The authors are grateful to the numerous owners who took time to fill the questionnaire, to the administrators of social media who shared the calls for participation in the survey, and to Thierry Legou for his intensive reading of this manuscript.

## Supporting information

**S1 Fig. Screenshot of the English version of the online questionnaire.**

**S2 Fig. Pictures of 55 sensory impaired and 33 sensory normal dogs from the present study illustrating the most typical coat colour patterns.** Pictures are sorted by group (HNVI, HIVN, HIVI, HNVN). Pictures of sensory impaired dogs with lesser white in the coat are framed in red. Pictures of sensory normal dogs with excess white coat are framed in green.

**S1 Video. Compulsive behaviours of two sensory impaired dogs filmed before and after behavioural adjustments by owners to the sensory impairments of their dogs.** The “initial” compulsive behaviours of these two dogs had been foremost considered as neurological signs prior to medical screening.

